# Disulfide bridge-dependent dimerization triggers FGF2 membrane translocation into the extracellular space

**DOI:** 10.1101/2023.04.12.536539

**Authors:** F Lolicato, JP Steringer, R Saleppico, D Beyer, J Fernandez-Sobaberas, S Unger, S Klein, P Riegerová, S Wegehingel, HM Müller, XJ Schmitt, S Kaptan, C Freund, M Hof, R Šachl, P Chlanda, I Vattulainen, W Nickel

## Abstract

Fibroblast Growth Factor 2 (FGF2) exits cells by direct translocation across the plasma membrane, a type I pathway of unconventional protein secretion. This process is initiated by PI(4,5)P_2_-dependent formation of highly dynamic FGF2 oligomers at the inner plasma membrane leaflet, inducing the formation of lipidic membrane pores. Cell surface heparan sulfate chains linked to glypican-1 (GPC1) capture FGF2 at the outer plasma membrane leaflet, completing FGF2 membrane translocation into the extracellular space. While the basic steps of this pathway are well understood, the molecular mechanism by which FGF2 oligomerizes on membrane surfaces remains unclear. In the current study, we demonstrate the initial step of this process to depend on C95-C95 disulfide-bridge-mediated FGF2 dimerization on membrane surfaces, producing the building blocks for higher FGF2 oligomers that drive the formation of membrane pores. We find FGF2 with a C95A substitution to be defective in oligomerization, pore formation, and membrane translocation. Consistently, we demonstrate a C95A variant of FGF2 to be characterized by a severe secretion phenotype. By contrast, while also important for efficient FGF2 secretion from cells, a second cysteine residue on the molecular surface of FGF2 (C77) is not involved in FGF2 oligomerization. Rather, we find C77 to be part of the protein-protein interaction interface through which FGF2 binds to the α1 subunit of the Na,K-ATPase, the landing platform for FGF2 at the inner plasma membrane leaflet. Using cross-linking mass spectrometry, atomistic molecular dynamics simulations combined with a machine learning analysis and cryo-electron tomography, we provide insights into a FGF2 dimerization interface that brings C95 residues in close proximity, resulting in disulfide bridged FGF2 dimers. We propose a mechanism by which they bind with high avidity to PI(4,5)P_2_ on membrane surfaces. We further propose a tight coupling between FGF2 secretion and the formation of ternary signaling complexes on cell surfaces, hypothesizing that C95-C95 bridged FGF2 dimers are functioning as the molecular units triggering autocrine and paracrine FGF2 signaling.

## Introduction

Beyond the classical ER/Golgi-dependent secretory pathway, multiple alternative mechanisms of protein secretion from cells have been discovered, processes collectively referred to as ‘unconventional protein secretion’ (UPS) (Rabouille, 2017; Dimou and Nickel, 2018; Pallotta and Nickel, 2020; Sparn et al., 2022b). The group of cargo proteins secreted by ER/Golgi-independent pathways is dominated by factors with fundamental functions in physiological processes such as inflammation and angiogenesis, among others, processes that are frequently linked to disease (Akl et al., 2016; Sitia and Rubartelli, 2018). Prominent examples for cargo proteins making use of unconventional mechanisms of protein secretion are Interleukin 1β (IL1β) and Fibroblast Growth Factor 2 (FGF2) (Nickel and Seedorf, 2008; Nickel and Rabouille, 2009; Zhang and Schekman, 2013; Rabouille, 2017; Sparn et al., 2022b). With regard to soluble cytoplasmic proteins lacking N-terminal signal peptides, two principal pathways have been identified. One of them is characterized by direct protein translocation across the plasma membrane, a process termed UPS Type I with FGF2 (Sparn et al., 2022b), HIV-Tat (Rayne et al., 2010; Schatz et al., 2018), Tau (Katsinelos et al., 2018; Merezhko et al., 2018; Katsinelos et al., 2021) and homeoproteins (Amblard et al., 2020; Joliot and Prochiantz, 2022) being examples. A second pathway mediating unconventional secretion of soluble cytoplasmic cargo proteins known as UPS Type III is based on intracellular vesicle intermediates such as autophagosomes or endocytic compartments (Malhotra, 2013; Zhang and Schekman, 2013; Rabouille, 2017; Dimou and Nickel, 2018; Ye, 2018; Liu et al., 2020). Interestingly, certain cargo proteins such as IL1β can be secreted via both type I and type III UPS pathways, depending on cell types and the physiological conditions that apply (Dupont et al., 2011; Liu et al., 2016; Evavold et al., 2017; Heilig et al., 2017; Monteleone et al., 2018; Chiritoiu et al., 2019; Liu et al., 2020; Pallotta and Nickel, 2020; Zhang et al., 2020). These examples reveal the complexity of secretory processes in mammalian cells as being much more diverse than previously assumed.

The unconventional secretory pathway of FGF2 is based on a small number of components, all of which are physically associated with the plasma membrane (Sparn et al., 2022b). The machinery can be classified into auxiliary factors and core machinery components. Auxiliary components are the Na,K-ATPase (Florkiewicz et al., 1998; Dahl et al., 2000; Ebert et al., 2010; Zacherl et al., 2015) and Tec kinase that binds to PI(3,4,5)P_3_ at the inner plasma membrane leaflet (Ebert et al., 2010; Steringer et al., 2012; La Venuta et al., 2016), factors that mediate initial steps of this pathway. The Na,K-ATPase has been demonstrated to be the first contact of FGF2 at the plasma membrane mediated by a direct physical interaction between its α1 subunit and FGF2 (Legrand et al., 2020). It is believed to serve as a landing platform of FGF2 at the inner plasma membrane leaflet, however, it has also been hypothesized to play an additional regulatory role in coupling FGF2 membrane translocation to the maintenance of the plasma membrane potential (Lolicato and Nickel, 2022; Sparn et al., 2022b). In addition, Tec kinase has been shown to make direct physical contact with FGF2, resulting in tyrosine phosphorylation of FGF2 at Y81, an interaction that is likely to occur downstream of the Na,K-ATPase (La Venuta et al., 2015; La Venuta et al., 2016; Lolicato and Nickel, 2022; Sparn et al., 2022b). This modification has been proposed to regulate the overall efficiency of FGF2 secretion, in particular in the context of cancer development (Ebert et al., 2010; Steringer et al., 2012; La Venuta et al., 2015; La Venuta et al., 2016; Sparn et al., 2022b).

Once FGF2 is handed over from the Na,K-ATPase and Tec kinase to the phosphoinositide PI(4,5)P_2_, the core mechanism of FGF2 membrane translocation into the extracellular space is triggered. It was demonstrated that the interaction with PI(4,5)P_2_ leads to FGF2 oligomerization on membrane surfaces (Steringer et al., 2012; Müller et al., 2015; Steringer et al., 2017), with hexamers being the most prominent oligomeric state linked to membrane pore formation (Steringer et al., 2017; Sachl et al., 2020). In addition to the high-affinity interaction of PI(4,5)P_2_ with a positively charged binding pocket containing K127, R128, and K133 (Temmerman et al., 2008; Temmerman and Nickel, 2009; Nickel, 2011; Steringer et al., 2012; Lolicato et al., 2022), molecular dynamics simulations suggest FGF2 to locally accumulate about 4-5 additional PI(4,5)P_2_ lipids through low-affinity interactions (Steringer et al., 2017). Thus, FGF2 hexamers may accumulate up to 30 PI(4,5)P_2_ lipids within a highly confined membrane surface area of about 10 nm^2^. At these sites, given the cone-shaped structure of PI(4,5)P_2_, local accumulation of PI(4,5)P_2_ will likely compromise the bilayer architecture and decrease the energy barrier for the formation of toroidal membrane pores (Lolicato and Nickel, 2022; Sparn et al., 2022b). Indeed, upon PI(4,5)P_2_-dependent FGF2 oligomerization on membrane surfaces, transbilayer diffusion of membrane lipids has been observed, suggesting a toroidal architecture of the membrane pores that were formed under these conditions (Steringer et al., 2012; Müller et al., 2015; Steringer et al., 2017; Dimou and Nickel, 2018; Pallotta and Nickel, 2020). With the heparan sulfate chains of GPC1 containing high-affinity binding sites for FGF2 in close proximity to the membrane surface, FGF2 oligomers have been shown to get captured and disassembled at the outer plasma membrane leaflet, completing FGF2 membrane translocation into the extracellular space (Zehe et al., 2006; Nickel, 2007; Sparn et al., 2022a). This step is unidirectional as heparan sulfate chains compete with high affinity against PI(4,5)P_2_ for the same binding site on the molecular surface of FGF2 (Steringer et al., 2017). Thus, in conclusion, the core machinery of FGF2 membrane translocation consists of PI(4,5)P_2_ and heparan sulfate chains linked to GPC1 on opposing sites of the plasma membrane. Along with this, FGF2 locally accumulates PI(4,5)P_2_ molecules through oligomerization concomitant with the formation of a membrane pore with a toroidal architecture (Pallotta and Nickel, 2020; Lolicato and Nickel, 2022; Sparn et al., 2022b).

The goal of the current study was to shed light on the structural principles that govern PI(4,5)P_2_-dependent oligomerization of FGF2 on membrane surfaces. Particular emphasis was given to the role of intermolecular disulfide bridges known to be present in PI(4,5)P_2_-triggered FGF2 oligomers. Furthermore, on a functional basis, it was known that a variant form of FGF2 lacking both of the two surface cysteine residues C77 and C95 is incapable of both oligomerization, membrane pore formation and secretion from cells (Müller et al., 2015; Steringer et al., 2017; Dimou and Nickel, 2018; Steringer and Nickel, 2018). However, the way these cysteines contribute to disulfide bridge formation and their specific roles in a cellular context remained unknown. In the current study, through a combination of biochemical, cell biological and structural techniques including cryo-electron tomography, cross-linking mass spectrometry, atom-scale biomolecular simulations and deep learning, we mechanistically dissected the functional roles of C77 and C95 in the sequence of events that constitute the unconventional secretory pathway of FGF2. As opposed to previous concepts, we found C77 not to be involved in FGF2 oligomerization. Rather, along with K54 and K60 (Legrand et al., 2020), we revealed a role for C77 in building the molecular interface through which FGF2 binds to the α1 subunit of the Na,K-ATPase, the starting point of this unusual pathway of protein secretion. By contrast, through the formation of disulfide bridges, we found C95 to be essential for PI(4,5)P_2_-dependent FGF2 oligomerization on membrane surface, producing the dimeric building blocks for higher FGF2 oligomers that can trigger the formation of lipidic membrane pores. Our findings further imply that the heparan sulfate chains of GPC1 containing high affinity binding sites for FGF2 to reverse this process by disassembling membrane pore forming FGF2 oligomers into C95-C95 disulfide-bridged dimers. The latter are likely to represent the primary ligands for the formation of FGF2 signaling complexes for autocrine and paracrine FGF signal transmission into cells (Decker et al., 2016; Nawrocka et al., 2020). We, therefore, propose unconventional secretion of FGF2 from cells and autocrine signal transmission into cells to represent tightly coupled processes.

## Results

### Cysteine residues on the molecular surface of FGF2 are required for efficient secretion of FGF2

To elucidate the precise mechanisms that turn C77 and C95 into critical *cis* elements required for unconventional secretion of FGF2 in a cellular context, we analyzed their individual roles in both FGF2 recruitment at the inner plasma membrane leaflet and FGF2 membrane translocation into the extracellular space (Fig. 1). Using a single molecule TIRF assay established previously (Dimou et al., 2019; Legrand et al., 2020; Lolicato et al., 2022), we found FGF2 mutants lacking either C77, C95 or both C77 and C95 to be impaired in membrane recruitment at the inner leaflet of the plasma membrane (Fig. 1A, 1B). This phenomenon was found to be statistically significant for C95A and C77/95A mutants of FGF2, indicating that oligomerization is required for robust FGF2 membrane recruitment at the inner plasma membrane leaflet. However, a moderate but consistently observed reduction of membrane recruitment was also observed for C77A. A potential reason for this subtle phenotype might be that this substitution causes a small disturbance of the interface between FGF2 and the α1 subunit of the Na,K-ATPase. To quantitatively assess FGF2 translocation to cell surfaces with the same set of FGF2 variants, we used a well-established cell surface biotinylation assay (Seelenmeyer et al., 2005; Zehe et al., 2006; Müller et al., 2015). Additionally, we also tested FGF2 variants in which cysteines were substituted with serine to exclude the observed phenotypes to depend on alanine substitutions of cysteines. As shown in Fig. 1C and 1D, a C77A or C77S substitution had a mild phenotype that was found to be statistically signifcant. By contrast, the substitution of C95 to alanine or serine in FGF2 caused a severe secretion phenotype (Fig. 1C and 1D). Of note, a combination of C77A (or C77S) and C95A (or C95S) further limited FGF2 secretion to background levels in a highly significant manner, suggesting that both C77 and C95 play important but most likely different roles in unconventional secretion of FGF2 from cells.

**Fig. 1:**
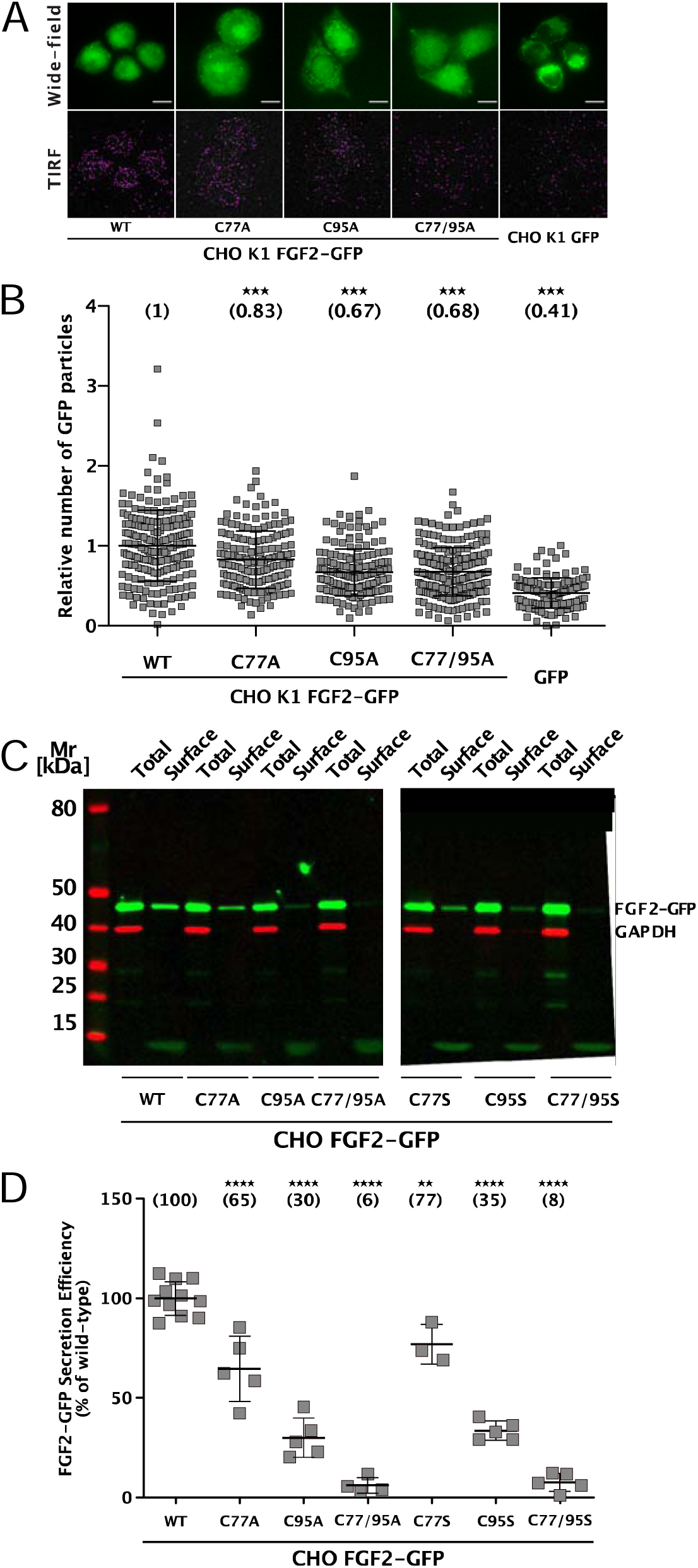
Cysteine residues in positions 77 and 95 of FGF2 play a role in its unconventional secretion from cells. (**A**) Representative wide-field and TIRF images of real-time single molecule TIRF recruitment assay conducted on stable CHO K1 cell lines overexpressing either wild-type (WT) or mutant (C77A, C95A, C77/95A) FGF2-GFP in a doxycycline-dependent manner. Beyond cell lines expressing various forms of FGF2-GFP, a GFP-expressing cell line was used to subtract GFP background. Wide-field images show the overall FGF2-GFP (or GFP) expression levels. Single FGF2-GFP (or GFP) particles recruited at the inner plasma membrane leaflet and detected within the TIRF field are highlighted with a pink circle. (**B**) Quantification of real-time single molecule TIRF recruitment assay conducted on the cell lines shown in panel A. Recruitment efficiency at the inner plasma membrane leaflet of FGF2-GFP wild-type was set to 1. Each square represents a single cell. Mean recruitment efficiency values are shown in brackets. Data are shown as mean with standard deviations (n = 4). Statistical analysis was based on a one-way ANOVA test performed in Prism (version 9.4.1), *** P ≤ 0.001. (**C**) Representative Western Blot of cell surface biotinylation assay conducted on stable CHO cell lines overexpressing either wild-type (WT) or mutant (C77A, C95A, C77/95A, C77S, C95S, C77/95S) FGF2-GFP in a doxycycline-dependent manner. Total cellular proteins and biotinylated surface proteins were analyzed. The analysis was conducted against GFP, to detect the various FGF2-GFP mutant forms, and GAPDH, both as a loading and a cellular integrity control. (**D**) Quantification of cell surface biotinylation assay conducted on the cell lines shown in panel D. Secretion efficiency of FGF2-GFP wild-type was set to 100%. Mean secretion efficiency values for each cell line are shown in brackets. Data are shown as mean with standard deviations (n = 4). Statistical analysis was based on a one-way ANOVA test performed in Prism (version 9.4.1), not significant (ns) P > 0.05, *** P ≤ 0.001).

### C95 is essential for PI(4,5)P_2_-dependent formation of FGF2 oligomers

To study the potential roles of C77 and C95 in PI(4,5)P_2_-dependent FGF2 oligomerization, we conducted both *in vitro* reconstitution experiments with purified components (Fig. 2) and FGF2 crosslinking experiments in cells (Fig. 3). For the first approach, we used giant unilamellar vesicles (GUVs) and purified variant forms of FGF2 as GFP fusion proteins (Fig. 2B) to quantify FGF2 oligomerization states by fluorescence correlation spectroscopy (FCS)/brightness analyses [Fig. 2A; (Steringer et al., 2017; Sachl et al., 2020); for details see Materials and Methods]. To allow for a systematic comparison with previous studies (Steringer et al., 2012; Müller et al., 2015; Steringer et al., 2017), phosphomimetic versions of FGF2 (Y81pCMF) were used under all experimental conditions. Consistent with earlier findings (Steringer et al., 2017), the wild-type version of FGF2-GFP was characterized by an average oligomeric state of about 6 to 8 subunits (Fig. 2A). By contrast, a FGF2 variant form lacking both C77 and C95 failed to oligomerize, an observation that again was consistent with previous findings (Müller et al., 2015; Steringer et al., 2017). Intriguingly, in the continued presence of C77, substituting C95 by alanine severely impaired PI(4,5)P_2_-dependent FGF2 oligomerization, an effect that was similar to what was observed with a FGF2 C77/C95A double substitution. By contrast, replacing C77 with alanine in the continued presence of C95 did not impact PI(4,5)P_2_-dependent FGF2 oligomerization to a significant extent (Fig. 2A).

**Fig. 2:**
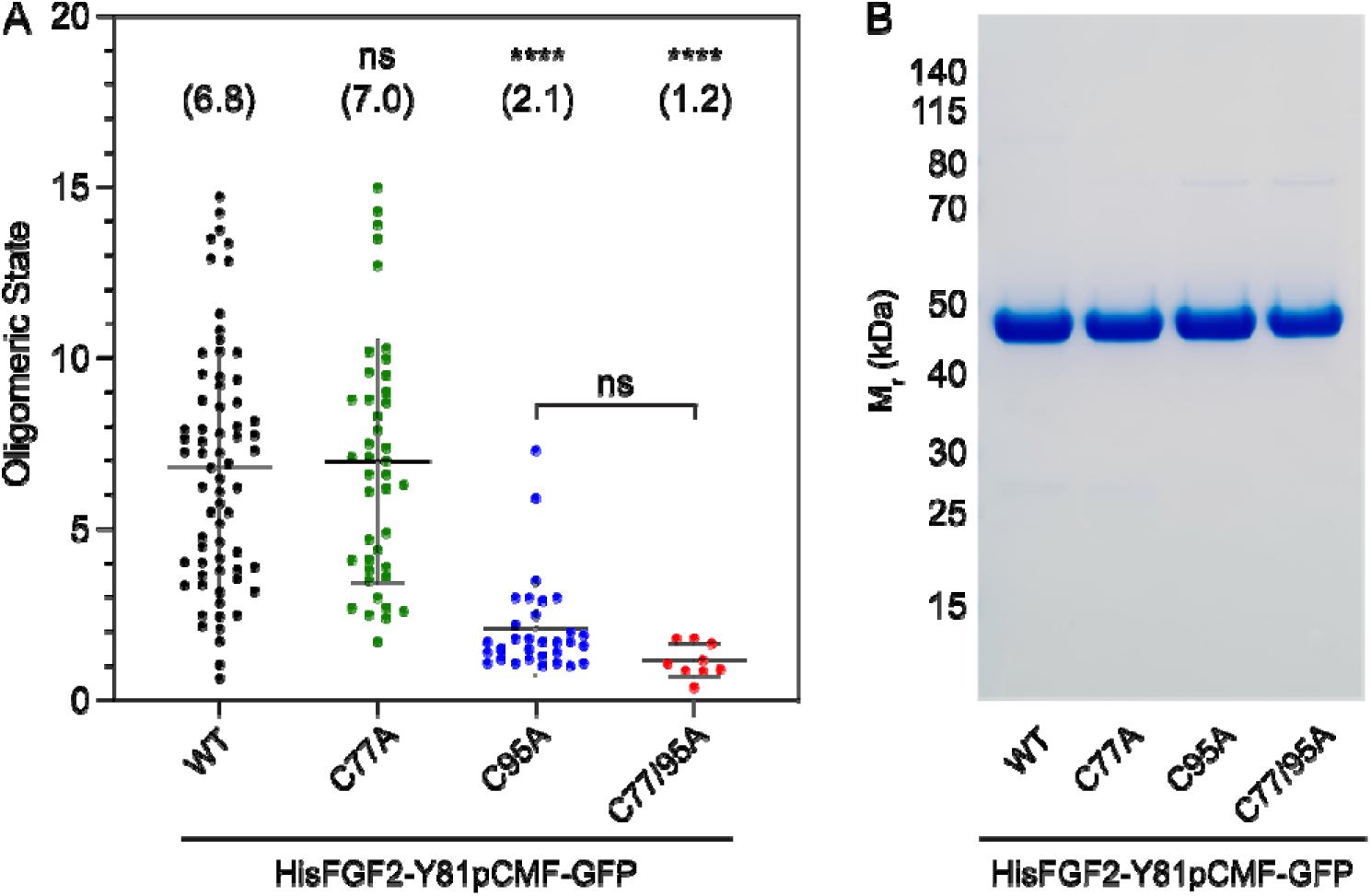
Formation of higher FGF2 oligomers on the membrane surface of GUVs depends on C95. **(A)** Oligomeric size distribution of FGF2-GFP variants. Giant unilamellar vesicles (GUVs) with a plasma membrane-like lipid composition containing 2 mol% PI(4,5)P_2_ were incubated with variant forms of His-tagged FGF2-Y81pCMF-GFP as indicated. The oligomer size was determined by brightness analysis as described in detail in Materials and Methods. Each dot corresponds to a data-point measured on a single GUV with the number of GUVs analyzed *n* (WT; *n* = 68, C77A; *n* = 42, C95A; *n* = 31, C77/95A; *n* = 9). Mean values with standard deviations are shown. One-way ANOVA with Tukey’ post hoc test was performed in Prism (version 9.4.1). Mean values are shown in brackets, not significant (ns) P >0.05, **** P ≤ 0.0001. Data distribution was assumed to be normal, but this was not formally tested. **(B)** SDS-PAGE analysis of FGF2-Y81pCMF-GFP variant forms indicated. Purified proteins were analyzed for homogeneity using Coomassie staining.

**Fig. 3:**
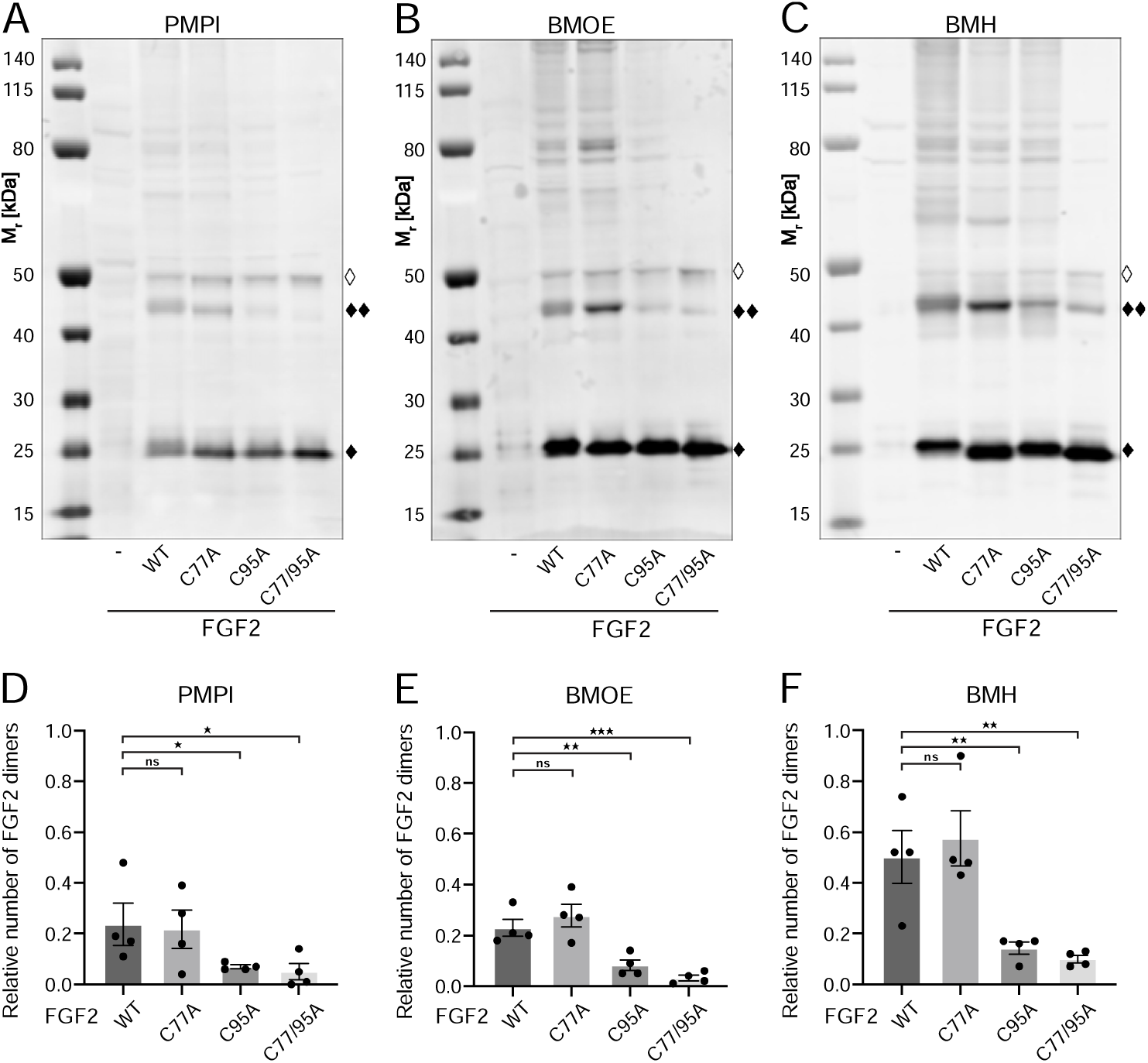
FGF2 dimer formation in cells depends on C95 as revealed by chemical cross-linking. Using cellular lysates, FGF2 dimer formation was analyzed by chemical cross-linking. The FGF2 variants (WT, C77A, C95A and C77/95A) were transiently expressed in HeLa S3 cells as constructs connecting the FGF2 open reading frame with GFP via a P2A site, producing stoichiometric amounts of untagged FGF2 and GFP, the latter used to label transfected cells. The corresponding cellular lysates were treated with three different crosslinkers: PMPI [N-p-maleimidophenylisocyanate; bifunctional crosslinker with a spacer length of 8.7 Å targeting sulfhydryl groups at one end (maleimide) and hydroxyl groups at the other end (isocyanate), Fig. 3A and D], BMOE [(bismaleimidoethane; bifunctional maleimide-based crosslinker with a short 8 Å spacer length targeting sulfhydryl groups), Fig.3B and E] or BMH [(bismaleimidohexane; bifunctional maleimide-based crosslinker with a long 13 Å spacer length targeting sulfhydryl groups), Fig.3C and F], respectively. Cross-linking products were analyzed by SDS-PAGE followed by Western blotting using polyclonal anti-FGF2 antibodies. **(A, B, C)** Representative examples of the Western analyses for each of the three crosslinkers described above. FGF2 monomers (18 kDa) are labeled with “♦”, FGF2 dimers (36 kDa) with “♦♦” and small amounts of monomeric full-length FGF2-P2A-GFP (∼50 kDa) with “◊”. **(D, E, F)** Quantification of FGF2 dimer to FGF2 monomer ratios. Signal intensities were quantified using a LI-COR Odyssey CLx imaging system. The FGF2 dimer to monomer ratios were determined in four independent experiments with the standard error of the mean shown, not significant (ns) P >0.05, * P ≤ 0.05, **≤ 0.01, *** P ≤ 0.001. Statistical analyses were based on two-tailed, unpaired t test using GraphPad Prism (version 9.4). Data distribution was assumed to be normal, but this was not formally tested.

To challenge these findings in a cellular context, we conducted cross-linking experiments in cellular lysates (Fig. 3). These studies focused on FGF2 dimers that have previously been shown to be abundantly present at the inner plasma membrane leaflet (Dimou et al., 2019). To systematically compare the wild-type form of FGF2 with the variant forms used in *in vitro* experiments (Fig. 2A and 2B), we transiently expressed FGF2 wt, C77A, C95A, and C77/95A in HeLa S3 cells. We employed an FGF2-P2A-GFP construct, leveraging the ‘self-cleaving’ P2A peptide to yield stoichiometric production of untagged FGF2 and GFP. GFP was utilized to monitor transfection efficiency. We used three chemical cross-linkers: PMPI, BMOE, and BMH, characterized by different spacer lenghts and chemical functionalities (details in Materials and Methods). The rationale of these experiments was that bifunctional crosslinkers targeting thiols would link FGF2 subunits into dimers only when the functional cysteine residues are present, replacing the disulfide bridge during oligomerization. As shown for representative examples in Fig. 3A, 3B, and 3C as well as quantified in Fig. 3D, 3E, and 3F, the amounts of cross-linked FGF2 dimers in cellular lysates (labeled with ♦♦) were similar between FGF2 wt and FGF2 C77A. By contrast, both FGF2 C95A and FGF2 C77/95A were severely impaired in dimer formation. In particular, using BMOE (bismaleimidoethane) and BMH (bismaleimidohexane), bifunctional crosslinkers with different spacer lengths that target the thiols of cysteine side chains, efficient crosslinking of FGF2 dimers was observed for FGF2 wt and C77A whereas a strong reduction of crosslinked dimers was found for FGF2 C95A and C77/C95A. This provides direct evidence for a C95-C95 disulfide bridge formed in FGF2 dimerization at the inner plasma membrane leaflet of cells.

The combined findings from *in vitro* experiments (Fig. 2) and cell-based analyses (Fig. 3) provide compelling evidence for C77 not to be involved in PI(4,5)P_2_-dependent FGF2 oligomerization. Instead, consistent with previous studies (Müller et al., 2015; Steringer et al., 2017), these data suggest that membrane-associated FGF2 dimers are formed by homotypic disulfide bridges linking C95 side chains.

### Cysteine 95 in FGF2 is essential for PI(4,5)P_2_-dependent membrane pore formation

In previous studies, we showed that PI(4,5)P_2_-dependent FGF2 oligomerization triggers the formation of membrane pores (Steringer et al., 2012; Müller et al., 2015; Steringer et al., 2017). While we reported previously that this process involves the formation of intermolecular disulfide bridges with a C77/95A variant of FGF2 being inactive (Müller et al., 2015; Steringer et al., 2017), the way disulfide bridges are formed based on potential contributions from these two cysteine residues remained unclear. As shown in Fig. 4A, using large unilamellar vesicles and His-tagged versions of recombinant FGF2 variant forms (Fig. 4B) along with a well-characterized dequenching assay to monitor membrane integrity (Steringer et al., 2012; Müller et al., 2015), FGF2 wt and FGF2 C77A displayed similar activities efficiently forming membrane pores through which liposome-enclosed fluorophores escape and dilute into the surroundings generating a fluorescent signal based on dequenching. By contrast, similar to the C77/95A variant form of FGF2, substituting C95 with alanine caused a dramatic drop in the formation of membrane pores (Fig. 4A). These findings are consistent with the experiments shown in Figs. 1, 2, and 3, linking FGF2 secretion from cells to membrane pore formation, a process triggered by PI(4,5)P_2_-dependent FGF2 oligomerization involving C95-mediated formation of disulfide bridges.

**Fig. 4:**
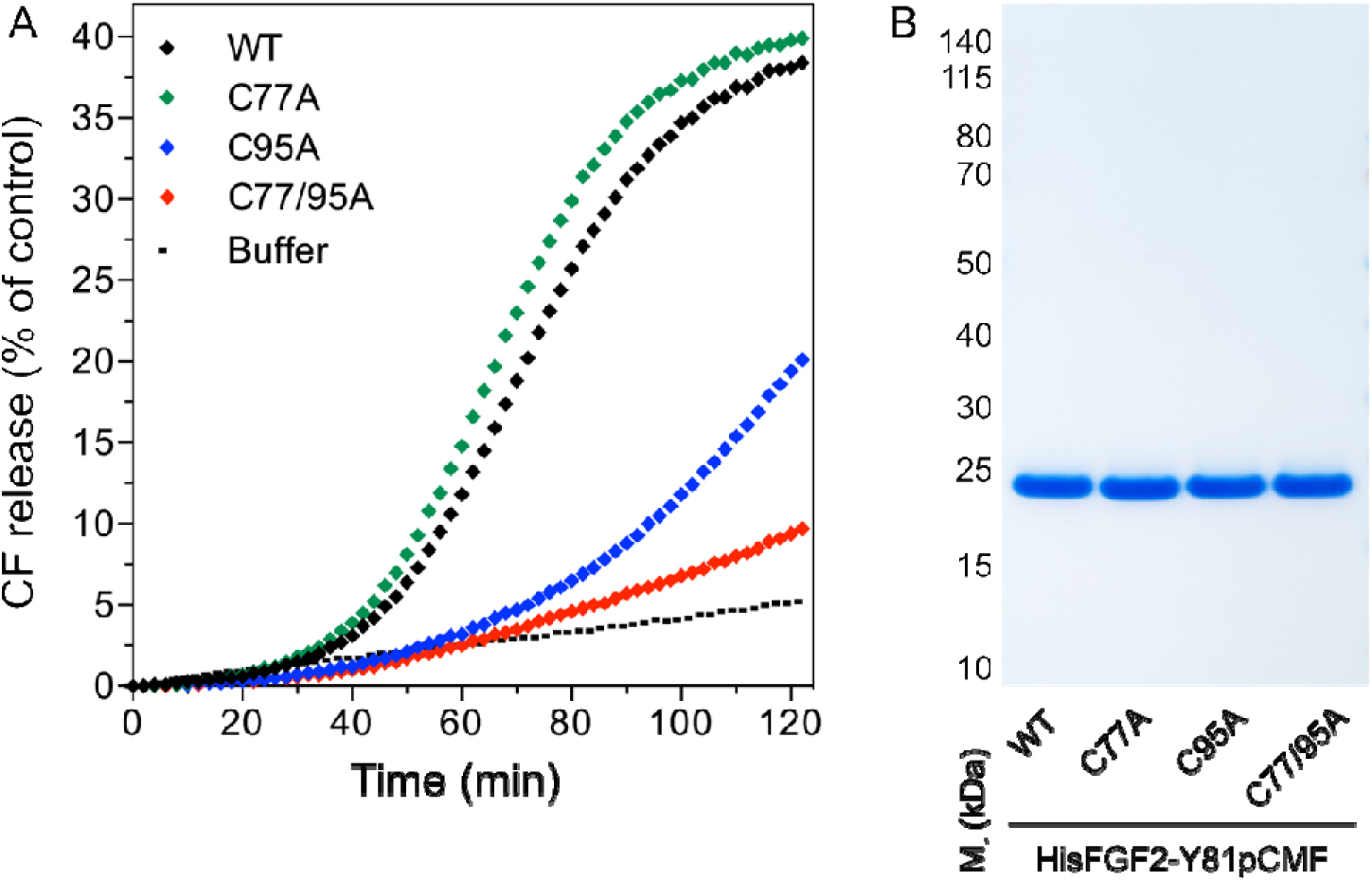
Membrane pore formation triggered by FGF2 oligomers depends on C95. Carboxyfluorescein was sequestered in large unilamellar liposomes containing a plasma-membrane-like lipid composition including 2 mol% PI(4,5)P_2_. **(A)** Liposomes were incubated with the His-tagged FGF2-Y81pCMF (2 µM)-based variant forms indicated (WT, C77A, C95A and C77/95A). Membrane pore formation was analyzed by measuring the release of luminal carboxyfluorescein quantified by fluorescence dequenching as detailed in Materials and Methods. The results shown are representative for three independent experiments. **(B)** Quality of the various recombinant proteins was analyzed by SDS PAGE and Coomassie staining.

### Cysteine 95 is essential for PI(4,5)P_2_-dependent FGF2 translocation across membranes

To analyze the role of C95 in a comprehensive manner through all steps of the unconventional mechanism of the FGF2 secretion pathway, we completed our *in vitro* studies by analyzing FGF2 translocation across the membrane of GUVs. These assays were based on previous work reconstituting the ability of FGF2 to physically traverse lipid bilayers based on an inside-out topology setup with PI(4,5)P_2_ and heparin (located in the lumen of GUVs and used as a surrogate of cell surface heparan sulfate chains) on opposing sides of GUV membranes (Steringer et al., 2017). His-tagged GFP fusion proteins of the various FGF2 forms described above (Fig. 2B) were tested for membrane recruitment, pore formation, and translocation into the lumen of GUVs. As shown in Fig. 5, GUVs were imaged in three independent fluorescence channels visualizing (i) FGF2 (GFP fluorescence), (ii) GUV lipid bilayers (rhodamine PE fluorescence) and (iii) an Alexa647 fluorophore used as a tracer to monitor membrane integrity. As shown in Fig. 5A for representative examples, radial intensity profiles were obtained to quantitatively compare fluorescence in the GUV lumen versus the exterior (Steringer et al., 2017). For FGF2 wt (Fig. 5A, sub-panel a) and FGF2 C77A (Fig. 5A, sub-panel b), a substantial increase in the GUV lumen could be observed, indicating FGF2 translocation across the membrane. By contrast, for FGF2 C95A (Fig. 5A, sub-panel c) and FGF2 C77/95A (Fig. 5A, sub-panel d), no difference between lumen and exterior could be observed, indicating a failure of FGF2 translocation. Under the conditions indicated, this experimental setup allowed for a statistical analysis of the formation of membrane pores (gray bars in Fig. 5B) along with luminal accumulation of FGF2-GFP within GUVs (green bars in Fig. 5B). All GUVs analyzed in Fig. 5 contained PI(4,5)P_2_ and, where indicated, luminal heparin. Based on the quantification shown in Fig. 5B and the representative images presented in Fig. 5C, the dynamic range of the experimental system was apparent from the comparison between GFP fusion proteins containing either FGF2 wt or FGF2 C77/95A. FGF2 wt caused membrane pore formation and, provided the presence of luminal heparin, translocated into the lumen of GUVs. By contrast, FGF2 C77/95A, while getting recruited to the surface of GUVs in a PI(4,5)P_2_-dependent manner, showed low activity with regard to both membrane pore formation and membrane translocation (Fig. 5A, 5B and 5C). Under the same conditions, when PI(4,5)P_2_ was substituted by a Ni-NTA lipid to mediate artificial membrane recruitment via the His tags of all GFP fusion proteins used in these experiments, both FGF2 wt and FGF2 C77/95A were low at background levels with regard to both membrane pore formation and membrane translocation (Fig. 6A, 6B and 6C). Based on this set of conditions, we analyzed the same parameters for FGF2 C77A and FGF2 C95A. Consistent with our findings documented in Figs. 2, 3, and 4, FGF2 C77A behaved similarly to FGF2 wt, efficiently forming membrane pores and, in the presence of luminal heparin, translocating across the membranes of GUVs containing PI(4,5)P_2_ (Fig. 5A, 5B and 5C). FGF2 C95A behaved differently, with significantly reduced activities regarding both membrane pore formation and membrane translocation (Fig. 5A, 5B and 5C). Like FGF2 wt and FGF2 C77/95A, both FGF2 C95A and C77A showed no activities when recruited via the Ni-NTA lipid on GUVs lacking PI(4,5)P_2_ (Fig. 6A, 6B and 6C). These findings are consistent with the data sets documented in Figs 2, 3, and 4, demonstrating the essential role of C95 in PI(4,5)P_2_-dependent FGF2 oligomerization concomitant with membrane pore formation and translocation across lipid bilayers.

**Fig. 5:**
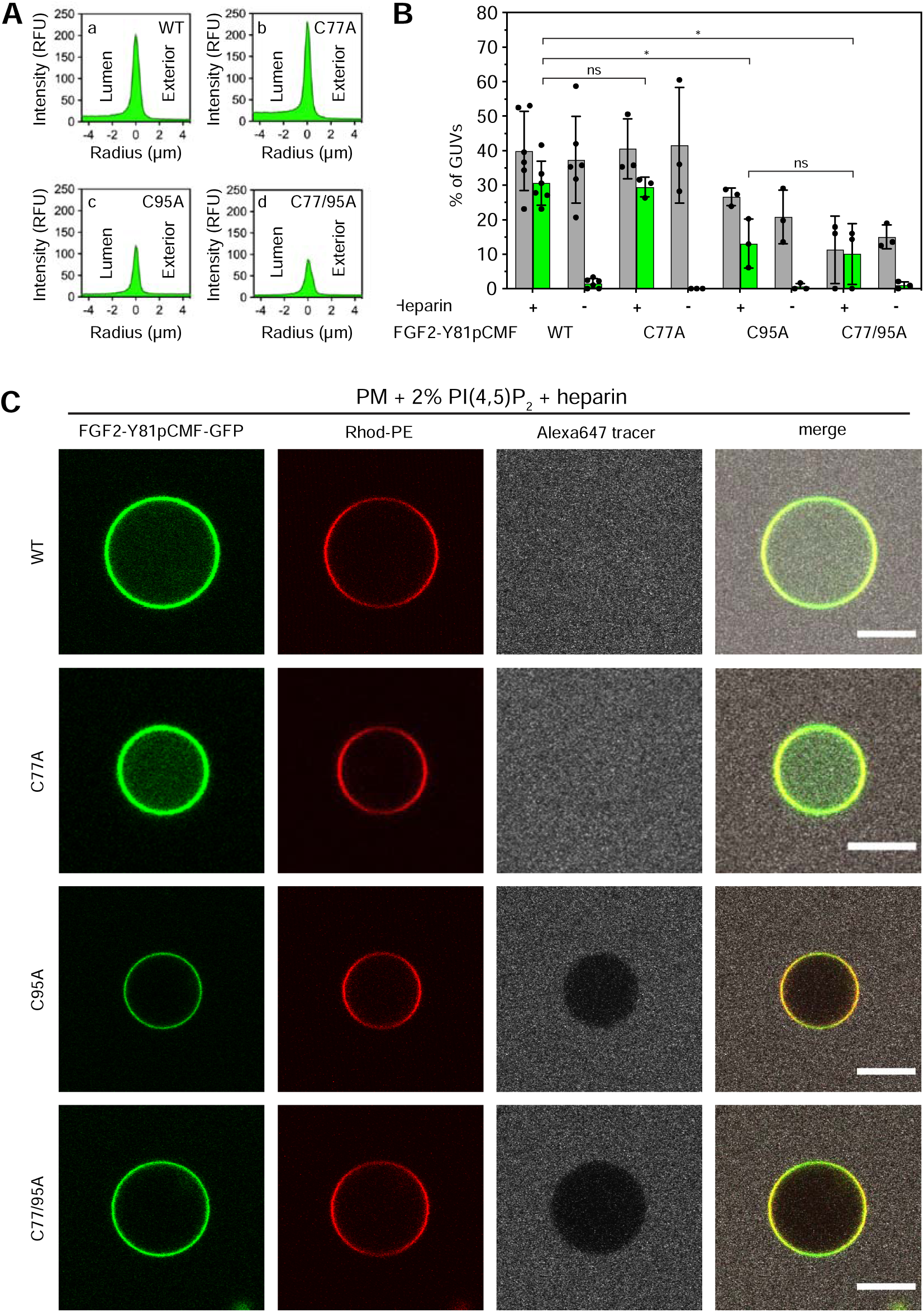
Full membrane translocation of FGF2 across GUV lipid bilayers depends on C95. Reconstitution of FGF2 membrane translocation with purified components. Giant unilamellar vesicles with a plasma membrane-like lipid composition containing 2 mol% PI(4,5)P_2_ were prepared in the presence or absence of long-chain heparins as described in detail in Materials and Methods. In brief, Rhodamine-PE (Rhod.-PE) was incorporated into the lipid bilayer during GUV preparation as membrane marker. After removal of excess heparin by low-speed centrifugation, GUVs were incubated with His-tagged FGF2-Y81pCMF-GFP (200 nM) variants as indicated and a small fluorescent tracer (Alexa647). Following 180 min of incubation luminal penetration of GUVs by FGF2-Y81pCMF-GFP and small tracer molecules was analyzed by confocal microscopy. **(A)** Radial intensity profiles of representative examples quantifying GFP fluorescence in the GUV lumen versus the exterior for FGF2-Y81pCMF-GFP wt (sub-panel a), C77A (sub-panel b), C95A (sub-panel c) and C77/95A (sub-panel d). **(B)** Quantification and statistical analysis of FGF2 membrane translocation and membrane pore formation. Gray bars indicate the percentage of GUVs with membrane pores with a ratio of Alexa647 tracer fluorescence in the lumen versus the exterior of ≥0.6. Green bars indicate the percentage of GUVs where membrane translocation of GFP-tagged proteins had occurred with a ratio of GFP fluorescence in the lumen versus the exterior of ≥1.6 being used as a threshold value. Each dot represents an independent experiment each of which involved the analysis of 20–120 GUVs per experimental condition. Mean values with standard deviations are shown. Statistical analyses are based on two-tailed, unpaired t-test performed in Prism (version 9.4.1), not significant (ns) P > 0.05, * P ≤ 0.05). Data distribution was assumed to be normal, but this was not formally tested. For details see Materials and Methods. **(C)** Representative confocal images of plasma membrane-like GUVs containing PI(4,5)P_2_ and long-chain heparins in the lumen after 180 min incubation with His-tagged FGF2-Y81pCMF-GFP (200 nM) variants as indicated and a small fluorescent tracer (Alexa647; scale bar = 10 µm).

**Fig. 6:**
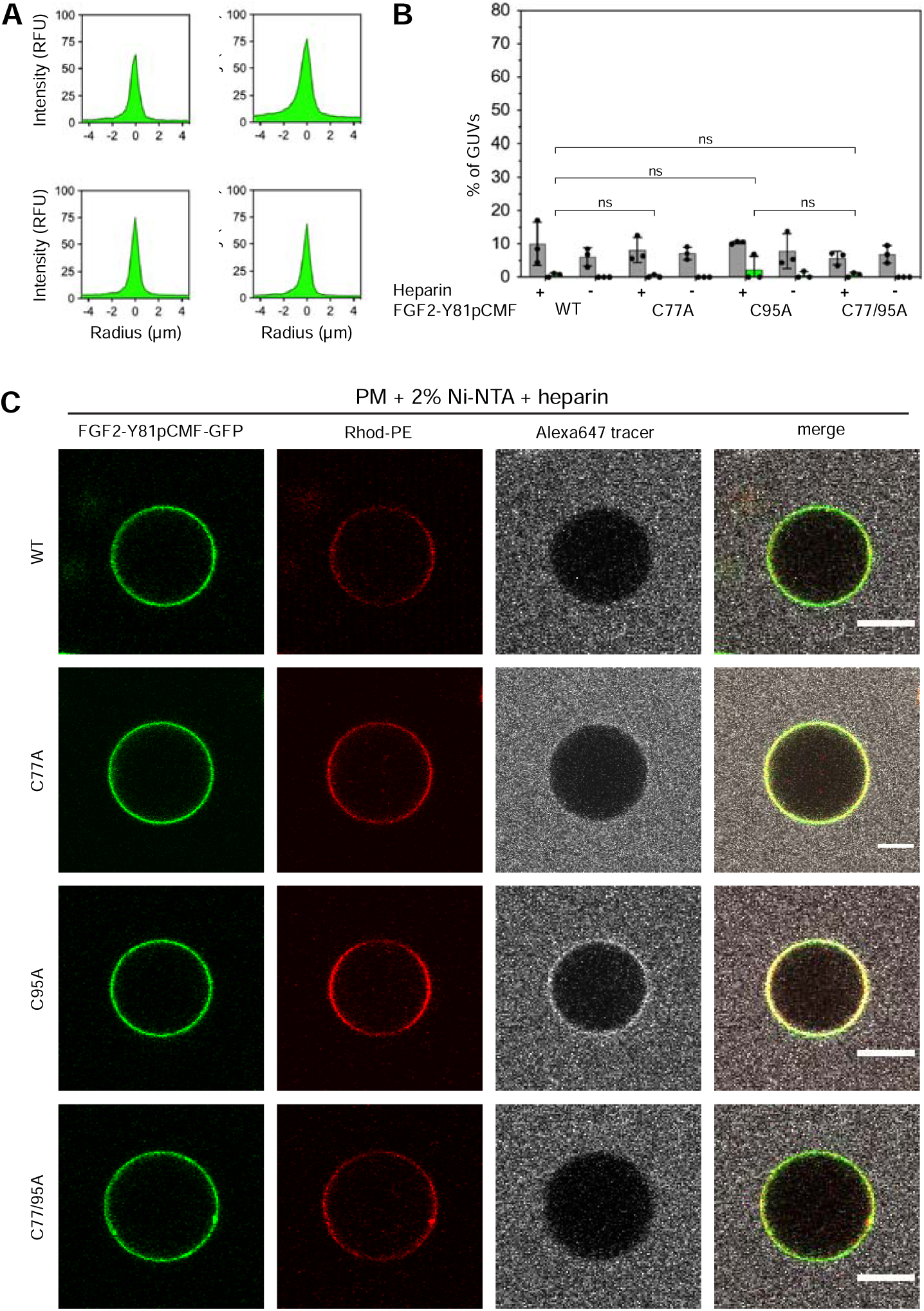
FGF2 membrane translocation across GUV lipid bilayers is abrogated when PI(4,5)P_2_ is substituted by a Ni-NTA lipid used to recruit His-tagged FGF2 fusion proteins. Giant unilamellar vesicles with a plasma membrane-like lipid composition containing 2 mol% Ni-NTA-lipid anchor were prepared in the presence or absence of long-chain heparins. Luminal penetration of GUVs by FGF2-Y81pCMF-GFP was analyzed by confocal microscopy as described in the legend to Fig.5. **(A)** Radial intensity profiles of representative examples quantifying GFP fluorescence in the GUV lumen versus the exterior for FGF2-Y81pCMF-GFP wt (sub-panel a), C77A (sub-panel b), C95A (sub-panel c) and C77/95A (sub-panel d). **(B)** Quantification and statistical analysis of FGF2 membrane translocation and membrane pore formation. Gray bars indicate the percentage of GUVs with membrane pores with a ratio of Alexa647 tracer fluorescence in the lumen versus the exterior of ≥0.6. Green bars indicate the percentage of GUVs where membrane translocation of GFP-tagged proteins had occurred with a ratio of GFP fluorescence in the lumen versus the exterior of ≥1.6 being used as a threshold value. Each dot represents an independent experiment each of which involved the analysis of 20–120 GUVs per experimental condition. Mean values with standard deviations are shown. Statistical analyses are based on two-tailed, unpaired t-test performed in Prism (version 9.4.1), not significant (ns) P > 0.05. Data distribution was assumed to be normal, but this was not formally tested. **(C)** Representative confocal images of plasma membrane-like GUVs containing Ni-NTA-lipid-anchor and long-chain heparins in the lumen after 180 min incubation with His-tagged FGF2-Y81pCMF-GFP (200 nM) variants as indicated and a small fluorescent tracer (Alexa647; scale bar = 10 µm).

### Cysteine 77 is a critical residue at the protein-protein interaction interface between FGF2 and the α1 subunit of the Na,K-ATPase

The experiments shown in Figs. 2 to 6 demonstrated C77 not to play any role in FGF2 oligomerization and membrane pore formation. Yet, substituting C77 by alanine caused moderate phenotypes in both FGF2 recruitment to the inner plasma membrane leaflet and FGF2 secretion from cells (Fig. 1B and 1D). In particular, a C77A substitution significantly enhanced the observed FGF2 secretion phenotype caused by a C95A substitution (Fig. 1D). When studying the position of C77 on the molecular surface of FGF2, it is evident that this residue is located in spatial proximity to two lysine residues (K54 and K60) that previously have been shown to be part of the protein-protein interface between FGF2 and the α1 subunit of the Na,K-ATPase [Fig. 7A; (Legrand et al., 2020)]. Therefore, in a cellular context, we hypothesized that a substitution of C77 by alanine may interfere with the initial recruitment step of FGF2 at the inner plasma membrane leaflet mediated by the α1 subunit of the Na,K-ATPase, a process that precedes PI(4,5)P_2_-dependent FGF2 oligomerization and membrane pore formation (Legrand et al., 2020). To test this possibility, as shown in Fig. 7, we used biolayer interferometry to study the binding kinetics of various forms of FGF2 towards the domain in the α1 subunit of the Na,K-ATPase to which FGF2 is known to bind [α1-subCD3; (Legrand et al., 2020)]. Both α1-subCD3 and the FGF2 variant forms indicated were expressed and purified to homogeneity as recombinant proteins (Fig. 7D). As detailed in Materials and Methods, α1-subCD3 was biotinylated and hooked up on optical sensors coated with streptavidin. To determine kinetic binding parameters between immobilized α1-subCD3 and FGF2, titration experiments were conducted with FGF2 wt concentrations ranging from 1 µM to 15 nM (Fig. 7B). These experiments revealed this interaction to be characterized by a K_D_ of 0.1 µM (± 0.01), an association constant k_on_of 1.46 x 10^4^ M^-1^ x s^-1^ (± 0.1 x 10^4^), and a k_off_constant of 1.46 x 10^-3^ s^-1^ (± 0.15 x 10^-3^). To compare the FGF2 variant forms indicated (FGF2 wt, C77A, C95A, C77/95A, K54/60E-C77A and K54/60E) with regard to their binding parameters towards α1-subCD3, all of them were used at a concentration of 1 µM, making use of the full dynamic range of the experimental system. As shown in Fig. 7C and quantified in Fig. 7E, FGF2 wt, and FGF2 C95A efficiently interacted with α1-subCD3. By contrast, FGF2 C77A was severely impaired in binding to α1-subCD3, a phenomenon shared with FGF2 K54/60E-C77A and K54/60E. These findings are consistent with previous observations demonstrating K54 and K60 to be part of the protein-protein interaction interface between FGF2 and α1-subCD3 (Legrand et al., 2020). Interestingly, FGF2 C77/95A was more severely impaired in interactions towards α1-subCD3 than FGF2 C77A, an observation that may indicate FGF2 dimerization via C95 to play a role in efficient interactions between FGF2 and the Na,K-ATPase This may also explain the finding that FGF2 C95A showed a slight but significant reduction in binding efficiency towards α1-subCD3 when compared with FGF2 wt (Fig. 7C and 7E). In conclusion, C77, along with K54 and K60, is part of the molecular surface of FGF2 that makes a direct physical contact with α1-subCD3. This, in turn, is consistent with C77A not being involved in PI(4,5)P_2_-dependent FGF2 oligomerization and membrane translocation, as demonstrated in the *in vitro* experiments shown in Figs. 2 to 6. Rather, cellular phenotypes observed for FGF2 C77A concerning recruitment at the inner plasma membrane leaflet and translocation into the extracellular space (Fig. 1) can be attributed to impaired binding efficiencies of FGF2 C77A towards the Na,K-ATPase (Fig. 7), the landing platform that mediates the initial contact of FGF2 with the plasma membrane as the starting point of its transport route into the extracellular space.

**Fig. 7:**
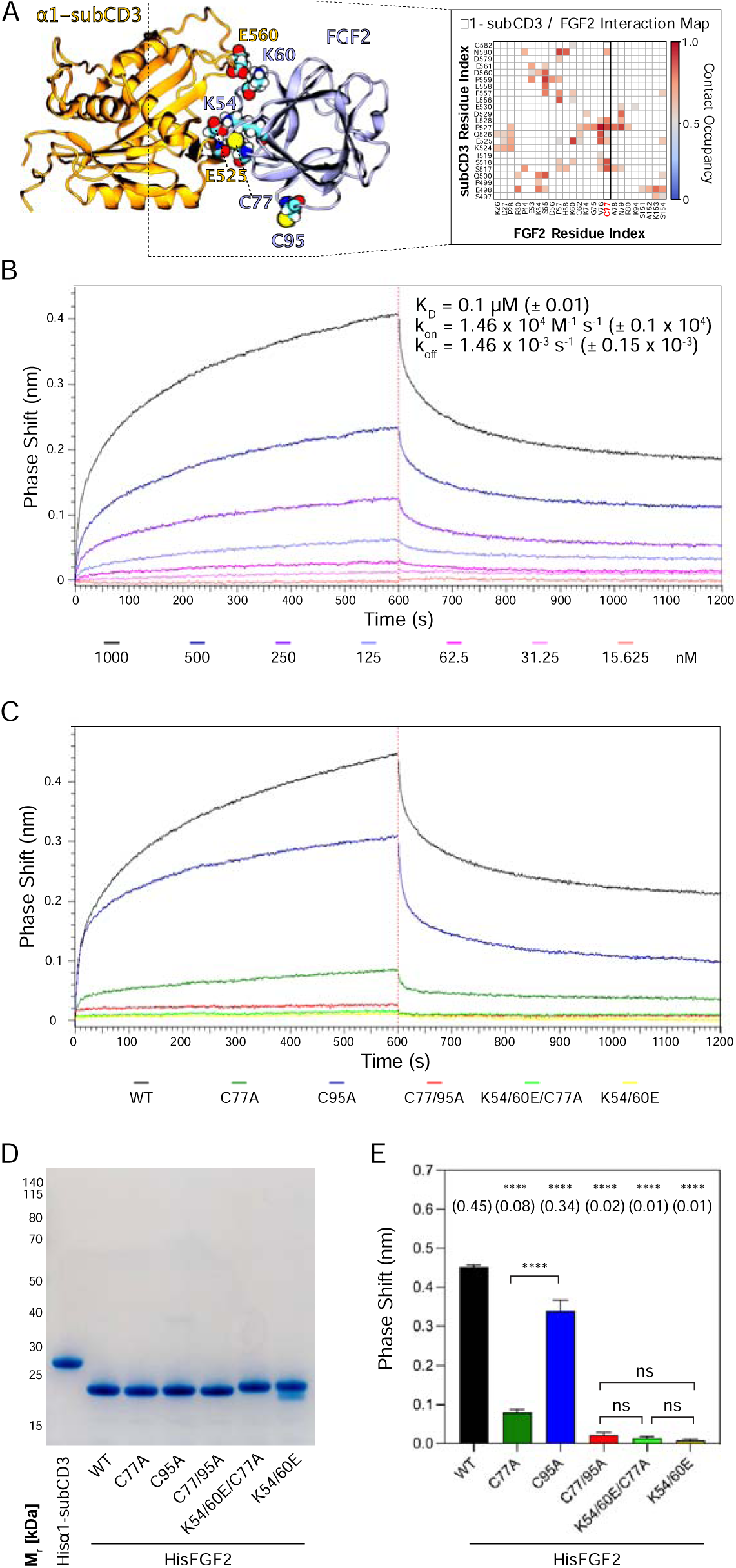
C77 is a component of the protein-protein interaction surface between FGF2 and the α1 subunit of the Na,K-ATPase. Kinetic analysis of the direct interaction of FGF2 with α1-subCD3 (Legrand et al., 2020). **(A)** FGF2 binds to α1 with K54, K60 and C77 being part of the protein-protein interaction interface [Figure adapted from (Legrand et al., 2020)]. **(B)** FGF2 directly binds to α1 in a dose-dependent manner. Biolayer interferometry (BLI) allows temporal resolution of association and dissociation. Biotinylated Hisα1-subCD3-WT protein was immobilised on Streptavidin sensors followed by incubation with His-tagged FGF2 wild type protein (HisFGF2-WT) at concentrations indicated. The data shown is representative of three independent experiments. Data were analyzed with Data Analysis HT 12.0 software (Sartorius) using a 1:1 binding model. See Material and Methods for details. Mean with standard deviations of K_D_, k_a_, and k_d_ values (n = 3) are given. **(C, D, E)** Comparison of FGF2 variants. **(C)** BLI measurements were conducted using immobilized Hisα1-subCD3 with FGF2 variants (1000 nM concentration) as indicated. The data shown is representative for four independent experiments. **(D)** The quality of HisFGF2 proteins was analyzed by SDS-PAGE and Coomassie staining. 3 µg of each variant were loaded as indicated. **(E)** Phase-shift at time point 600 s. Mean values with standard deviations of four independent experiments are shown. One-way ANOVA with Tukey’ post hoc test was performed in Prism (version 9.4.1). Mean values are shown in brackets, not significant (ns) P > 0.5, **** P ≤ 0.0001. Data distribution was assumed to be normal, but this was not formally tested.

### Simulations reveal that the C95-C95 interaction interface dominates the observed dimerization interfaces

The results from the biochemical and cell-based experiments shown in Figs. 1 to 7 provided direct evidence for a crucial role of C95 in FGF2 dimerization on membrane surfaces based on the formation of disulfide bridges. To investigate the likelihood of this interaction occurring independently of disulfide bond formation, we created 360 initial structures (see 360 Degree Analysis, Materials and Methods) in which two FGF2 monomers (not disulfide-bridged linked) attached to the membrane surface in close proximity underwent different orientations with respect to each other, degree by degree, and each of these systems was simulated for 0.5 microseconds. The goal was to find out all dimerization interfaces where C95 is involved. Visualization of the simulations immediately revealed that the C95 residues of the monomers sought proximity to each other. This was not observed for C77 residues (Fig. 8A). Analysis of the simulation data using a combination of dimensionality reduction (Fig. 8B) and clustering with machine learning techniques revealed that FGF2 dimers formed 8 noteworthy clusters (Fig. 8C). In the cluster with the largest population (Cluster 2, Fig. 8D), the C95-C95 residues of the two monomers were less than 1 nm apart (Fig. 8E). This spatial arrangement is stabilized by a network of salt bridges between the two monomers, in which the residues K85, E86, K118, E66, and D98 played a crucial role (Fig. 8F). Among the observed clusters, this cluster is the only one where the C95-C95 pair is compatible to disulfide bridge formation. The findings reveal that the FGF2 dimer structure depicted in Cluster 2 (Fig. 8F) forms spontaneously, with the C95-C95 residues being in close proximity and oriented at a specific distance conducive to disulfide bridge formation. This spontaneous configuration does not rely solely on forming the disulfide bridge, suggesting that the covalent bond formation plays a pivotal role in stabilizing the interface during the membrane translocation process.

**Fig. 8:**
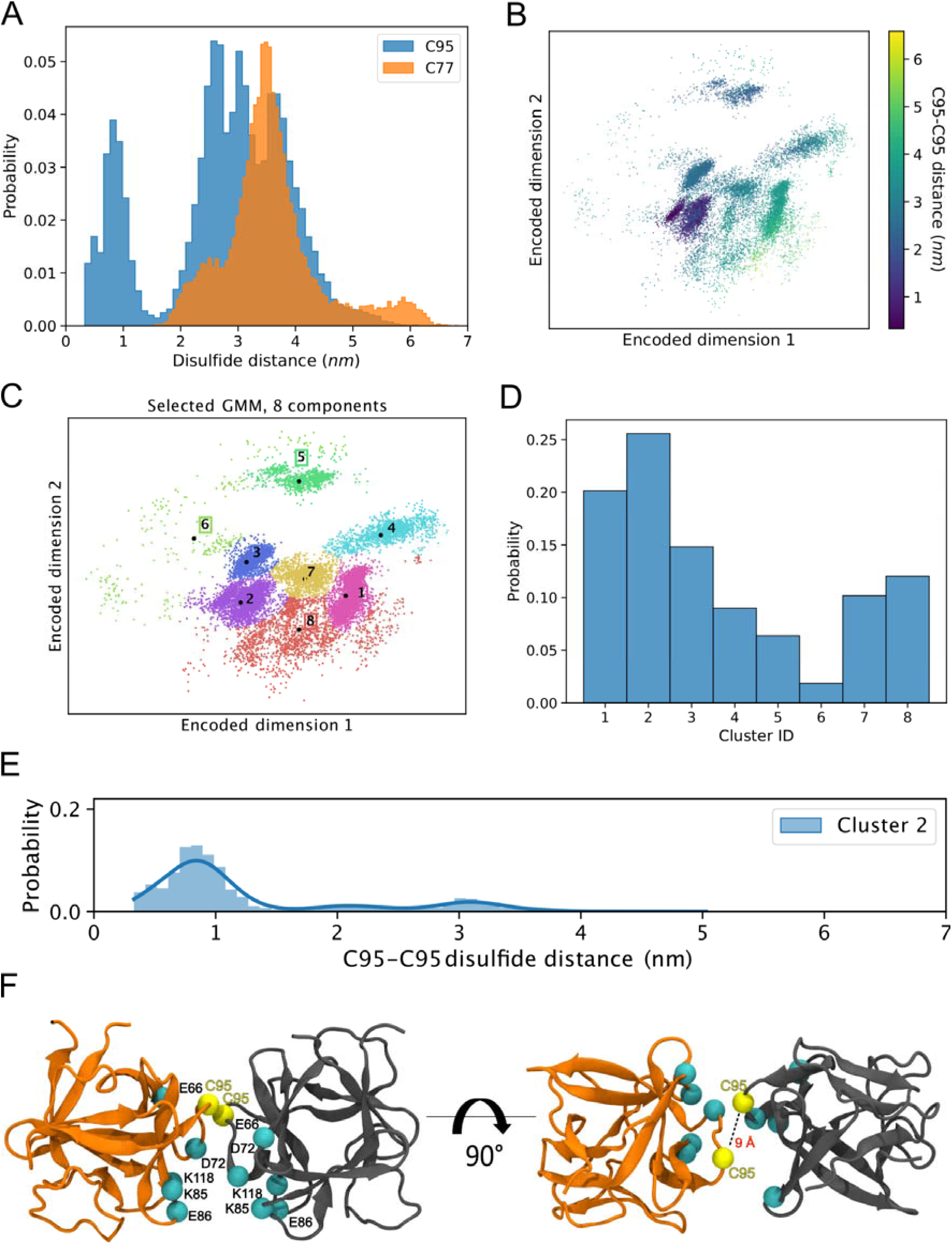
Simulations reveal that the C95-C95 interaction interface forms independently of the disulfide bridge. **(A)** The distribution of the C95-C95 and C77-C77 distance shows that only the former can come within 1 nm of each other during the unbiased 360 MD simulations, where one FGF2 monomer was systematically rotated to explore all possible C95-involved dimerization interfaces. **(B)** The C-alpha atoms of the two monomers were reduced to a 2D representation using an orthogonal autoencoder. The points are colored with the C95-C95 distance. A cluster with low C95-C95 distance is visible in the representation. **(C)** The encoded space was clustered with a Bayesian Gaussian Mixture Model (GMM) to find regions of distinct conformational structures. The 8 identified clusters are indicated in the figure, along with the cluster mean shown in black dots and the corresponding cluster label. **(D)** Populations of the individual clusters in the GMM. Cluster 2 have the largest population amongst the 8 identified clusters. **(E)** The C95-C95 distance indicates that the highest occupied cluster (Cluster 2) also has the highest likelihood of low C95-C95 distance (below 1 nm). **(F)** Representative structure of the FGF2 dimer from Cluster 2 showing C95 residues in proximity and crucial residues responsible for salt bridge interactions.

### Characterization of C95-C95 disulfide-bridged FGF2 dimers employing computational approaches

We continued the atomistic simulations further, aiming to obtain structural insights into this process. We utilized computational approaches to study in detail the protein-protein interface of C95-C95 bridged FGF2 dimers. We generated seven C95-C95 FGF2 dimerization interfaces using three different techniques. As explained in the ‘Materials and Methods’ section in detail, these approaches included atomistic molecular dynamics simulations performed previously (Steringer et al., 2017), the ROSETTA protein-protein docking protocol (Gray et al., 2003; Wang et al., 2005; Wang et al., 2007; Chaudhury and Gray, 2008) as well as predictions generated with the AlphaFold2-Multimer v3 package (Evans et al., 2021; Jumper et al., 2021). To stabilize the dimerization interface found in the initial structures, we conducted 1-µsec-long MD simulations in water with a C95-C95 disulfide linked FGF2 dimer as the starting point (Fig. 9A, subpanels a and b). These simulations produced a consistent interface in five independent simulations, confirming the presence of two ion pairs (E86 – K118 and E99 – K85) that had previously been suggested to play a role in FGF2 dimerization (Steringer et al., 2017). They further revealed that the interface is highly flexible, allowing the two monomers to rotate until they reach a stable conformation (Fig. 9A, subpanel b). This flexibility is likely to be critical in the context of FGF2-induced pore formation that requires a substantial remodeling of the lipid bilayer. For example, when FGF2 oligomers become accommodated inside toroidal membrane pores, a high degree of freedom is essential to maintain interactions of FGF2 with PI(4,5)P_2_ in the presence of high membrane curvature. The dimers’ final configuration was randomly placed in ten different orientations, 2 nm away from a POPC membrane surface containing 2 PI(4,5)P_2_ molecules (Fig. 9B, subpanel a). These configurations were then simulated for 1 µsec. The interaction between the dimer and the membrane preserved the original dimeric interface and revealed the interaction to occur in two steps (Fig. 9B, subpanel b-c). First, a single FGF2 molecule binds to a single PI(4,5)P_2_ molecule (Fig. 9B, subpanel b), followed by the second FGF2 molecule binding to the other PI(4,5)P_2_ molecule (Fig. 9B, subpanel c). The free energy profile (Fig. 9B, subpanel d) indicates that the final state in which both FGF2 subunits were attached to the membrane surface, is energetically strongly favorable with a free energy value of −30 kT. The free energy calculations were done using only two PI(4,5)P_2_ molecules, however, the dimer still had a free energy value comparable to the system where the monomer was bound to five PI(4,5)P_2_ molecules (Lolicato et al., 2022). These findings suggest that the C95 disulfide dimer with a higher avidity has a stronger affinity for the membrane than the FGF2 monomer, even under the conditions that were used for the MD simulations described above. This indicates that the dimer might be more likely to interact with low-abundance PI(4,5)P_2_ molecules in a cellular context. Additionally, it suggests that FGF2 dimerization might occur before PI(4,5)P_2_-dependent FGF2 binding to the membrane, possibly triggered by the interaction of FGF2 with the α1 subunit of the Na,K-ATPase, the initial contact of FGF2 with the inner plasma membrane leaflet.

**Fig. 9:**
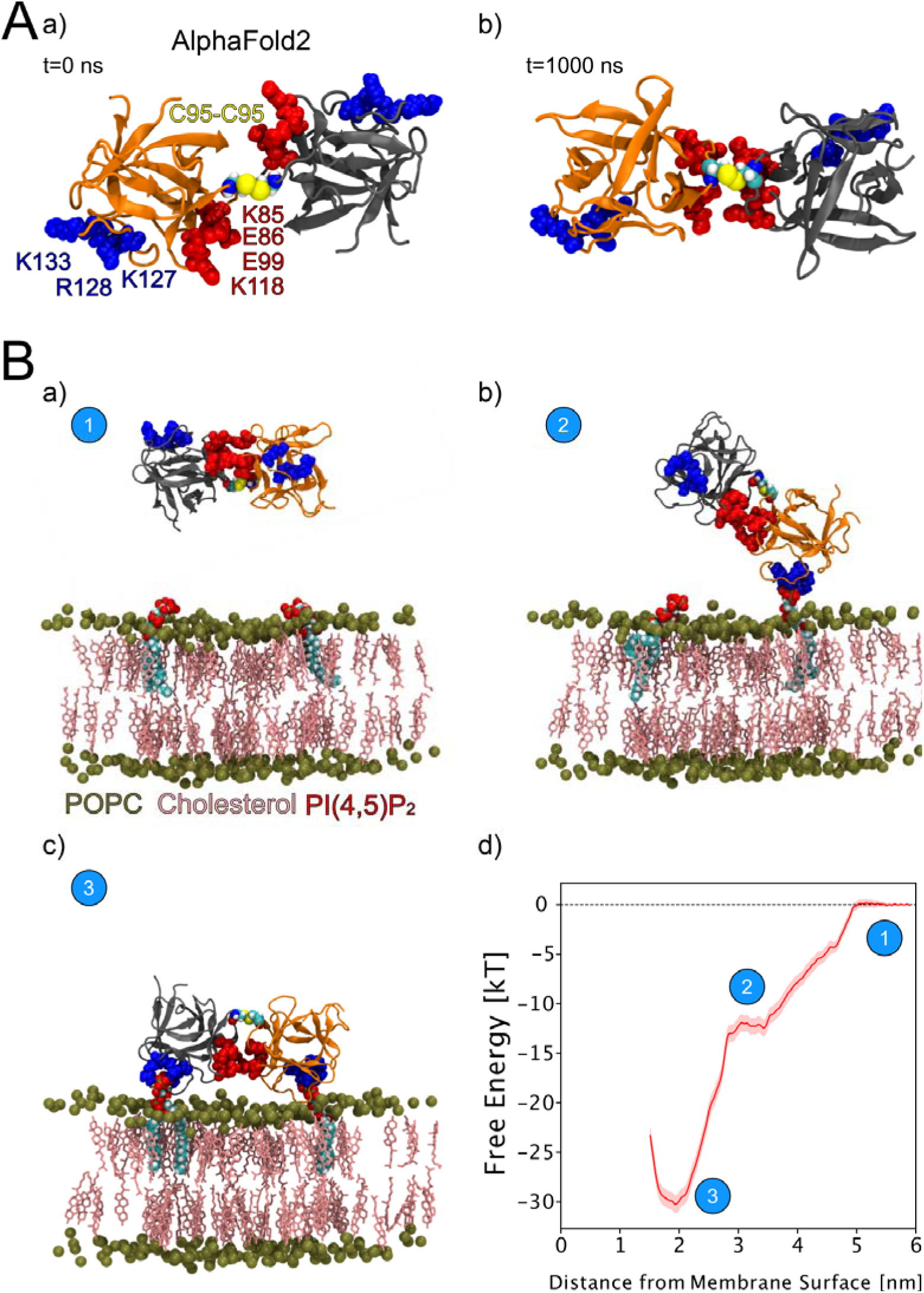
Characterization of C95 disulfide-bridged FGF2 via Molecular Dynamics Simulations. **(A)** The AlphaFold2 model dimeric interface’s stability (subpanel a) was tested by conducting 1-microsecond-long MD simulations in water, which revealed the interface’s high flexibility (subpanel b). **(B)** Unbiased all-atom MD simulations were used to sample the FGF2 dimer-membrane interaction pathway, mediated via the experimentally known PI(4,5)P_2_ binding pocket (K127, R128, K133). The free energy profile of FGF2 dimer-membrane interaction was determined from biased (umbrella sampling) MD simulations and plotted against the center of the mass distance of FGF2 dimer from phosphate atoms of the interacting membrane surface. Subpanels a-c show the interaction pathway’s initial, intermediate, and final states, while subpanel d shows the free energy profile.

### Characterization of FGF2 dimer interface employing cross-linking mass spectrometry

To further analyze the spatial arrangement of the interface that mediates FGF2 dimerization on membrane surfaces, we employed cross-linking mass spectrometry (XL-MS, Fig. 10). A bifunctional crosslinker targeting amino groups in the side chains of amino acids (disuccinimidyldibutyric urea; DSBU) was used to cross-link FGF2 dimers on liposomal surfaces. Following enzymatic cleavage using the Lys-C protease (see ‘Materials and Methods’ for details), the resulting peptides were subjected to a mass spectrometric analysis. To focus on inter-molecular cross-links in the protein-protein interface of membrane-bound FGF2 dimers, we exclusively considered distances produced from identical residues in the subunits of FGF2 dimers. Experiments were conducted with FGF2 wt and FGF2 C77/95A in the presence and absence of PI(4,5)P_2_-containing liposomes. In Fig. 10A, under the experimental conditions indicated, cross-linked peptides are provided by the homotypic pairs of amino acid residues they were derived from. Consistent with previous experiments demonstrating FGF2 oligomerization to depend on membrane surfaces (Steringer et al., 2012; Müller et al., 2015; Steringer et al., 2017), cross-linked peptides were more abundantly found in the presence of PI(4,5)P_2_-containing liposomes. A structural analysis revealed the collection of cross-linked peptides to be compatible with a FGF2 dimerization interface that brings C95 residues into proximity, enabling the subsequent formation of a disulfide bridge. Specifically, the observed cross-links involving K74-K74, K85-K85 and K94-K94 support an interface that brings C95 residues from two FGF2 molecules into direct contact. These findings are supportive of the molecular dynamics simulations of the membrane-bound FGF2 dimer as shown in Fig. 10B, with the Cα-Cα distances of these pairs measuring below the 26.4 Å that are given as DSBU cross-linking distance in rigid molecules (Iacobucci et al., 2019; Piersimoni and Sinz, 2020). Intriguingly, cross-linked peptides could also be observed with the C77/95A variant form of FGF2, suggesting a protein-protein interface whose formation does not depend on disulfide formation. However, once C95 residues are brought into proximity in FGF2 wt molecules disulfide formation can take place and further stabilize the interface. The data from the XL-MS experiments further suggest a second independent dimerization interface that may play a role in the formation of higher FGF2 oligomers in which C95-C95-bridged FGF2 dimers are used as building blocks (see average size of FGF2 oligomers to be hexamers as shown in Fig. 2). This interface is characterized by cross-links involving K34-K34, K54-K54, K60-K60 and K143-K143. and is compatible with a spatial arrangement in which both PI(4,5)P_2_ binding pockets are pointing to the membrane surface (Fig. 10C). In conclusion, using an independent experimental approach, the data shown in Fig. 10 directly support the functional experimental data of this study (Figs. 1 to 7) and molecular dynamics simulations (Figs. 8 and 9), pointing at a FGF2 dimerization interface that brings C95 residues in close proximity that is compatible with the formation of a disulfide bridge.

**Fig. 10:**
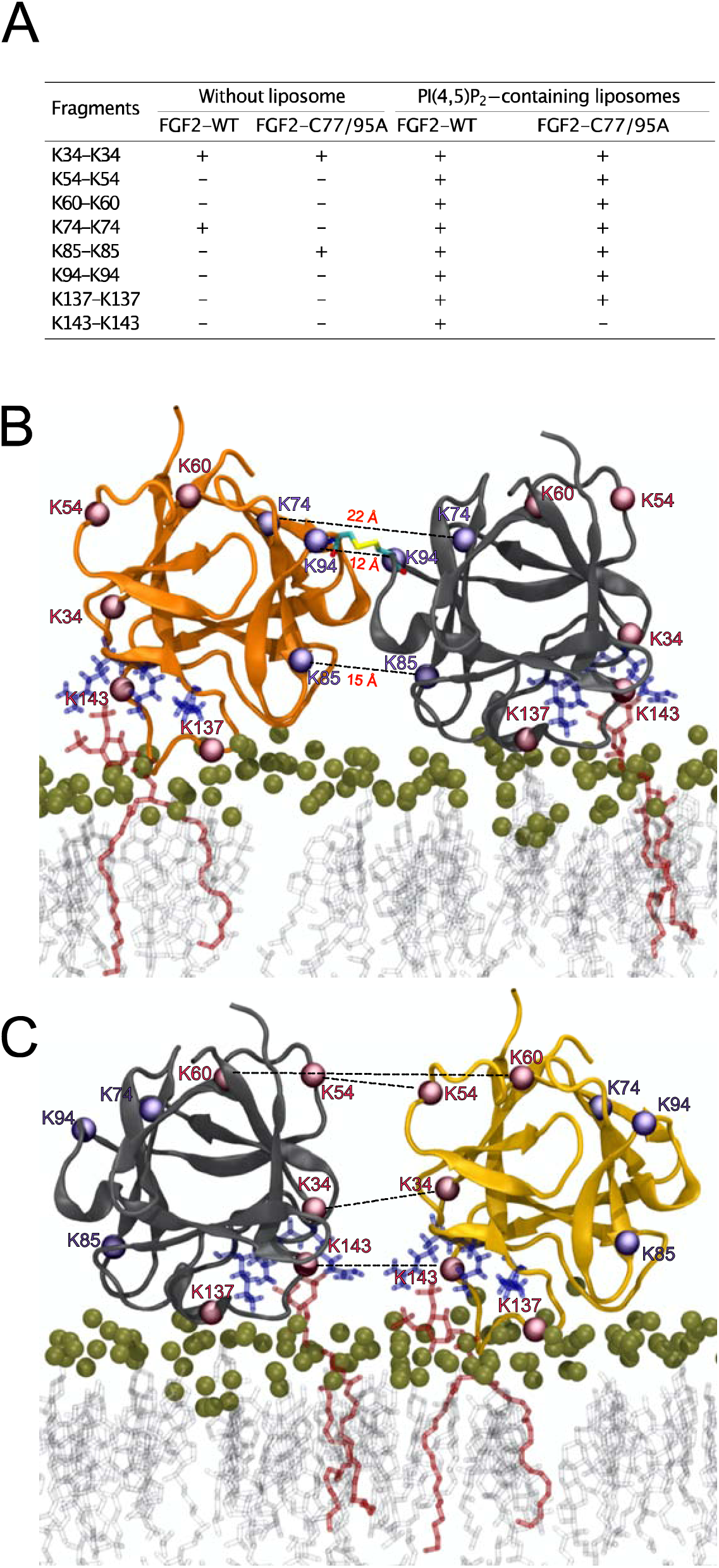
Cross-Linking Mass Spectrometry visualization of FGF2 dimer interfaces. **(A)** Overview of the inter-cross-linked fragments for His-tagged FGF2-WT and FGF2-C77/95A in the presence and absence of PI(4,5)P_2_-containing liposomes. **(B)** Liposome-induced cross-linking findings align seamlessly with molecular dynamics simulations’ C95-dependent dimer model interface. Fragments K74-K74, K85-K85, and K94-K94 can be positioned within the simulation model at a Cα-Cα distance below 23 Å. This concurs with the theoretical maximum Cα-Cα distance of approximately 26.4 Å for DSBU-linked lysine residues. **(C)** Cross-linking mass spectrometry data are compatible with an additional dimerization interface. It is incompatible with disulfide bridge formation but consistent with a membrane-bound FGF2 dimer configuration. Fragments K34-K34, K54-K54, K60-K60, and K143-K143 can be positioned at a Cα-Cα distance below 23 Å.

### Visualization of membrane-associated FGF2 dimers by cryo-electron tomography

To visualize interactions of FGF2 with membrane surfaces at a molecular scale, we conducted cryo-electron tomography (cryo-ET) using FGF2 bound to liposomes in a PI(4,5)P_2_-dependent manner. Since FGF2 has a low molecular weight (18 kDa) and thus is challenging to be imaged by cryo-ET, we took advantage of the fact that FGF2 fusion proteins are functional in both cell-based assays (Fig. 1) and *in vitro* reconstitution experiments (Figs. 2 and 5). Since the Halo tag (33 kDa) with its strongly acidic isoelectric point decreases liposome tethering (Lolicato et al., 2022), we used a His-FGF2-Y81pCMF-Halo fusion protein with a molecular weight of approximately 51 kDa. FGF2-Y81pCMF-Halo (10 µM) was incubated with large unilamellar vesicles [LUVs; 2 mM lipids with a plasma membrane-like composition containing PI(4,5)P_2_; (Temmerman et al., 2008; Temmerman and Nickel, 2009; Steringer et al., 2012; Müller et al., 2015; Steringer et al., 2017)] for four hours at 25°C. Proteoliposomes were vitrified by plunge freezing into liquid ethane (Vitrobot, ThermoFischer Scientific) as described in ‘Materials and Methods’. Tilt series were acquired at Krios-GIF-K2 either in-focus using a Volta phase plate (VPP) or at nominal defocus (−3 µm) without the VPP. Cryo-electron tomograms revealed small densities of His-FGF2-Y81pCMF-Halo bound to the membrane surface of LUVs (Fig. 11A, subpanels a-c). Based on the number of visible Halo domains contained in single particles, our analysis revealed monomers (e.g. Fig. 11A, subpanel d), dimers (e.g. Fig. 11A, subpanel e) and higher oligomers (e.g. Fig. 11A, subpanel f) of His-FGF2-Y81pCMF-Halo. In the latter case, Halo tags could be observed on both sides of the membrane, suggesting that higher His-FGF2-Y81pCMF-Halo oligomers are capable of spanning the lipid bilayer (Fig.11A, subpanel f).

**Fig. 11:**
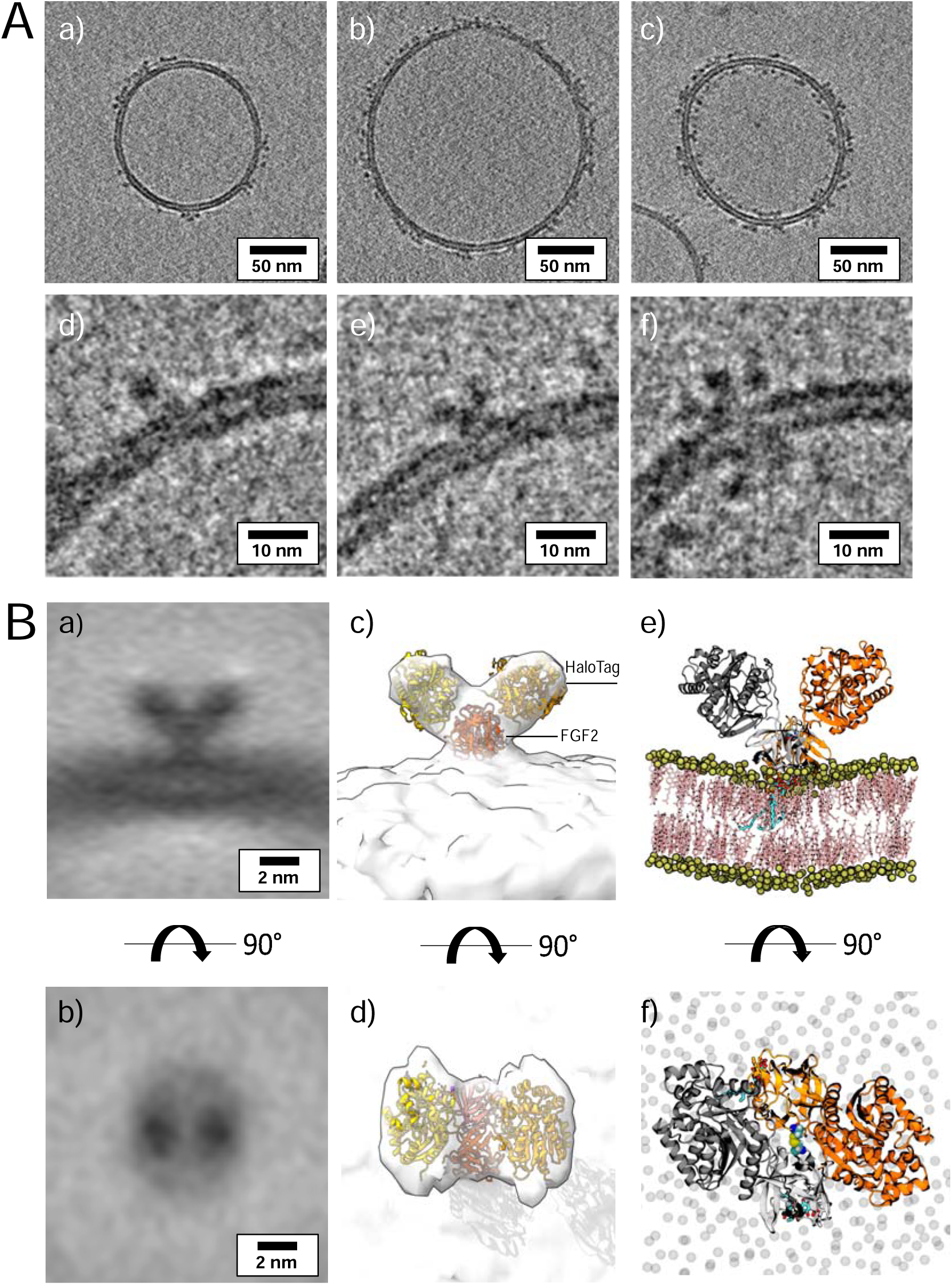
Cryo-electron tomography visualization of FGF2 dimer in proteo-liposomes. **(A)** Example slices of cryo-electron tomograms (sub-panels a-c; 2.3 nm thickness) showing His-FGF2-Y81pCMF-Halo bound to PI(4,5)P_2_-containing liposomes acquired at nominal defocus −3 µm. Magnified views from FGF2-Halo monomers, dimers and higher oligomers (sub-panels e-f). **(B)** Subtomogram average of the V-shaped FGF2-Halo dimer interacting with the membrane of PI(4,5)P_2_-containing liposomes (sub-panels a-b). Subtomogram average of “V-shaped” FGF2 dimers were manually picked using a dipole model in Dynamo (number of particles = 186). Top and side views are shown in subpanel a and b, respectively; c-d) 3D map with manually fitted crystal structures of two Halo domains (PDB:4KAJ) and two FGF2 (PDB:1BFF) monomers; e-f) Atom-scale molecular dynamics simulation model of V-shape C95 disulfide bridged FGF2-Halo dimer stable over 500 nanoseconds.

Interestingly, dimeric forms of His-FGF2-Y81pCMF-Halo bound to PI(4,5)P_2_-containing liposomes appeared as V-shaped structures (Fig. 11A, subpanel e). This finding prompted us to analyze these dimers in more detail, using subtomogram averaging of subvolumes extracted from cryo-electron tomograms obtained with VPP (Fig. 11B). A total of 184 V-shaped particles, which showed a similar structure to the one presented in Fig. 11A, were picked manually using a dipole model and subjected to subtomogram averaging workflow using Dynamo [(Castano-Diez et al., 2012; Castano-Diez, 2017; Castano-Diez et al., 2017; Navarro et al., 2018); for details, see ‘Materials and Methods’]. The resulting 2-fold symmetrized subtomogram average was visualized as a 3D volume (Fig. 11B; subpanels c-d) in ChimeraX (Pettersen et al., 2021). Using the known crystal structures of the Halo domain (PDB:4KAJ) and the FGF2 monomer (PDB:1BFF), we were able to manually fit two FGF2 monomers to the membrane proximal region and two Halo domains protruding away from the membrane surface into the subtomogram average. This indicates that the V-shaped particle corresponds to a FGF2-Y81pCMF-Halo dimer where FGF2 is interacting with the membrane surface of the liposome. However, due to a limited number of particles and heterogeneity in oligomeric states, the resolution of the average is limited and not sufficient to provide direct information about the FGF2 dimerization interface. Therefore, we used AlphaFold2-Multimer (v3) to generate FGF2-HALO dimers that fit into the 3D density in an accurate manner. Remarkably, four of the top-ranked structures resembled a V-shape FGF2-Halo dimer. However, among these structures, only the fourth-ranked one contained the experimentally known PI(4,5)P_2_ binding pocket in an orientation that is compatible with PI(4,5)P_2_-dependent membrane binding. Intriguingly, the C95 residues from the two FGF2 subunits were found in the interaction interface (Fig. 12A and 12B). Using this structure as a starting point, we substituted the predicted FGF2 dimer with the one obtained from the MD simulations (Figs. 8 and 9). In addition, AlphaFold2-Multimer (v3) predicted the positions of the two Halo domains in a way that was not compatible with the electron densities obtained from cryo-ET (Fig. 12A and 12B). Therefore, we performed a rigid body fit using ChimeraX software to place the single Halo domains into the density map, using the AlphaFold-Multimer structure as a template. Finally, the modeled V-shape dimer was positioned on the surface of a model membrane and subjected to a 500 nanosecond simulation. As shown in Figure 11–video supplement 1, the dimer was stable over the simulated time period. As depicted in Fig. 11B (subpanels e-f), the resulting FGF2-Halo dimer is based on a consistent data set combining cryo-ET data, structural predictions from AlphaFold2-Multimer (v3) and MD simulations. The data provide an initial structural understanding of how FGF2 dimerizes on membrane surfaces. The proposed dimerization interface is compatible with the functional data from both cell-based studies and in vitro experiments presented in this study.

**Fig. 12:**
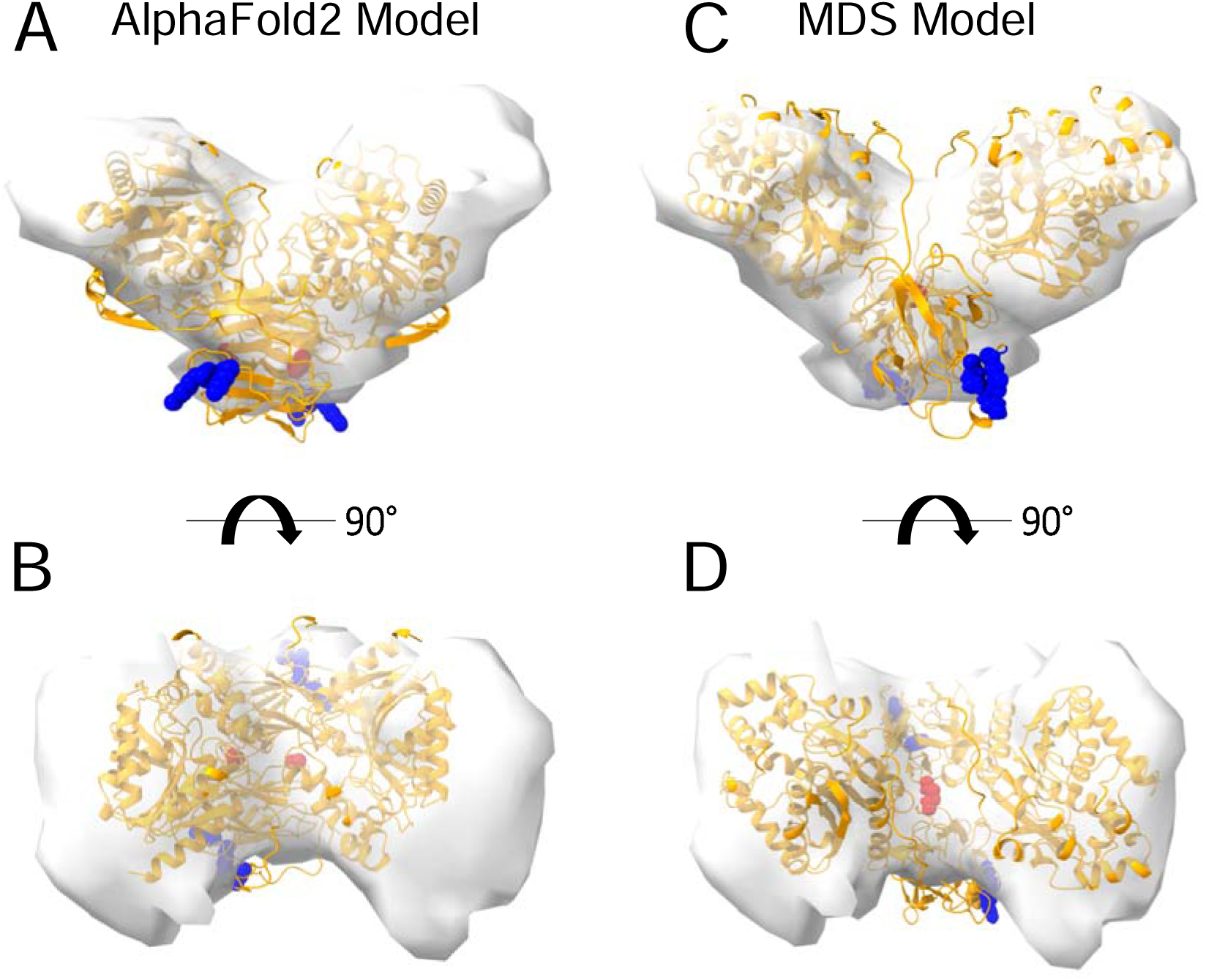
Cryo-electron tomography visualization of FGF2 dimer in proteo-liposomes. From a top and side view perspective, a comparison between the AlphaFold2 Multimer v3 model (panels **A-B**) and the one used for molecular dynamics simulations (panels **C-D**). The AlphaFold model accurately predicted the orientation of FGF2 dimer, with the PI(4,5)P_2_ binding residues correctly positioned for a membrane-bound state (blue residues). However, the two cysteines 95 (red residues), although located at the interface, were observed to be distant. To improve the model, we replaced the FGF2 dimer with the C95-C95 disulfide bridged dimer interface characterized with MD simulations (Fig. 9). Furthermore, we used the “Fit in Map” command provided by the ChimeraX software to locally optimize the fit of one of the two Halo domains’ atomic coordinates into the density map.

## Discussion

The principal machinery and basic aspects of the molecular mechanism by which FGF2 can physically traverse the plasma membrane to get access to the extracellular space have been revealed in great detail in recent years (Dimou and Nickel, 2018; Pallotta and Nickel, 2020; Sparn et al., 2022b). However, the mechanism by which FGF2 oligomerizes in a PI(4,5)P_2_-dependent manner on membrane surfaces concomitant with the formation of a lipidic membrane pore continued to be a mystery. In particular, the molecular events remained elusive by which the highly dynamic process of PI(4,5)P_2_-dependent FGF2 oligomerization triggers the remodeling of the rather stable plasma membrane lipid bilayer into a lipidic membrane pore with a toroidal architecture, the intermediate structure through which FGF2 oligomers can move towards the extracellular space. This process concludes in a GPC1-depending manner, a cell surface heparan sulfate proteoglycan that captures FGF2 oligomers as they penetrate PI(4,5)P_2_-dependent membrane pores, followed by disassembly into FGF2 signaling modules on cell surfaces (Zehe et al., 2006; Nickel, 2007; Sparn et al., 2022a). To address the long-term goal of a mechanistic understanding of how FGF2 oligomers can trigger the biogenesis of lipidic membrane pores in a PI(4,5)P_2_-dependent manner, it is crucial to understand how FGF2 molecules assemble into oligomers at the inner leaflet of plasma membranes.

In previous work, we identified two surface cysteines in position 77 and 95, respectively, that are fully conserved among all mammalian forms of FGF2 (Müller et al., 2015; Steringer et al., 2017). Of note, these cysteines cannot be found in FGF family members containing N-terminal signal peptides for ER/Golgi-dependent secretion, suggesting a specialized role in unconventional secretion of FGF2 (Steringer and Nickel, 2018). Manipulation of these cysteine residues by either NEM-mediated alkylation or substituion by alanines resulted in the inability of FGF2 to oligomerize on membrane surfaces in a PI(4,5)P_2_-dependent manner (Müller et al., 2015). Such conditions further caused a failure of FGF2 to form membrane pores, to translocate across membranes and to get secreted from cells (Müller et al., 2015; Steringer et al., 2017). When oligomeric species of FGF2 were analyzed by native gelelectrophoresis or by comparing migration behavior by reducing and non-reducing SDS-PAGE, evidence could be collected that intermolecular disulfide bridges play a role in PI(4,5)P_2_-dependent FGF2 oligomerization on membrane surfaces (Müller et al., 2015). However, it remained unclear as to whether both cysteines are involved and how disulfide bridges are arranged to form FGF2 dimers and higher oligomers.

With the current study, we now obtained key insights into the structural foundations of how FGF2 dimerizes on membrane surfaces. In conjunction with functional *in vitro* studies and cell-based experiments, this became possible only through a combination of cryo-ET, atomistic molecular dynamics simulations, machine learning analyses, structural predictions from AlphaFold2-Multimer and insights from cross-linking mass spec experiments, providing a data set suited to solve this challenging problem. In particular, we revealed a FGF2 dimerization interface that contains a disulfide bridge that depends on C95, one of two cysteine residues on the molecular surface of FGF2. Of note, disulfide bridge formation driving FGF2 oligomerization was found to be completely independent of C77, a second residue with a thiol side chain that is localized nearby C95 on the molecular surface of FGF2. Intriguingly, our data from machine learning analysis (Fig. 8), atomistic molecular dynamic simulations (Fig. 9), and cross-linking mass spectrometry (Fig. 10) indicate that this dimerization interface assembles independently of C95-C95 disulfide formation, meaning the stability of this configuration is not derived solely from the formation of the disulfide bridge. Instead, it offers kinetic stability essential for disulfide bridge formation at the inner leaflet of the plasma membrane, indicating that the formation of disulfide-bridged FGF2 dimers plays a critical role in stabilizing the dimerization interface during FGF2 membrane translocation. This, in turn, explains why C95 substitutions by alanine or serine cause severe FGF2 secretion phenotypes. These findings clarify the molecular configuration by which disulfide bridges are formed in membrane-associated FGF2 dimers. Furthermore, the inability of C77 to form disulfide bridges had important implications of the overall structure of higher FGF2 oligomers. In particular, we propose a second non-covalent FGF2-FGF2 interaction interface to exist, resulting in the formation of higher FGF2 oligomers in which disulfide bridges and non-covalent FGF2-FGF2 interaction interfaces occur in an alternating arrangement. The C95-dependent disulfide-mediated interface could be revealed in this study (Figs. 8, 9A, 9B, 10 and 11). The second disulfide-indendent interface is likely to be represented by the one identified in this study through cross-linking mass spectrometry experiments (Fig. 10A and 10C). The combination of these interfaces will be key for future studies, revealing the structure-function relationship of higher FGF2 oligomers forming lipidic membrane pores during FGF2 translocation into the extracellular space.

The limitation to just one type of disulfide bridge in FGF2 oligomers also has implications on how these oligomers are disassembled on cell surfaces, mediated by GPC1 (Sparn et al., 2022a; Sparn et al., 2022b). We propose the product of this process to be C95-C95 linked FGF2 dimers, generated by the GPC1-dependent disruption of the predicted second non-covalent FGF2-FGF2 interface. Of note, recombinant forms of C95-C95 FGF2 dimers stabilized by chemical crosslinking have been demonstrated to be efficient FGF2 signaling modules (Decker et al., 2016; Nawrocka et al., 2020). These observations led us to hypothesize that the unconventional secretory pathway of FGF2 and the generation of FGF2 signaling units are tightly coupled processes, with the dimeric FGF2 signaling modules that drive FGF2 signaling already forming at the inner plasma membrane leaflet. In this concept, FGF2 dimers transiently oligomerize into higher oligomers as intermediates of FGF2 membrane tranlocation, a process that has been shown to occur within a time interval of just 200 ms (Dimou et al., 2019). By the action of GPC1, we propose membrane-inserted FGF2 oligomers to be reconverted into FGF2 dimers at the outer leaflet, the principal untits of FGF2 signal transduction (Plotnikov et al., 1999; Plotnikov et al., 2000; Schlessinger et al., 2000).

Starting from the original observation of PI(4,5)P_2_-dependent oligomerization to involve intermolecular disulfide bridges (Müller et al., 2015), the current study resolved the amino acid code of this phenomenon, demonstrating an exclusive role for C95. Nevertheless, as revealed in this study, a substitution of C77 to alanine caused FGF2 secretion from cells to become less efficient (Fig. 1). This phenomenon could be explained with the proximity of C77 to K54 and K60 on the molecular surface of FGF2, residues that previously have been demonstrated to be part of the molecular interaction interface between FGF2 and the α1 subunit of the Na,K-ATPase (Legrand et al., 2020). Indeed, using biolayer interferometry to study the kinetics of FGF2/α1 interactions, we revealed a C77A substitution alone to severely weaken this interface. Triple substitutions (K54/60E-C77A) caused a complete failure of FGF2 to bind to the α1 subunit of the Na,K-ATPase. With these findings, we could resolve a long-standing question regarding the molecular mechanism of FGF2 membrane translocation, the differential roles of C77 and C95 with regard to FGF2 interactions with α1 and disulfide-mediated dimerization of FGF2 on membrane surfaces. Our observations have important implications for future studies revealing the mechanisms by which higher FGF2 oligomers can trigger and become accommodated in lipidic membrane pores, based on the flexible hinge produced by the C95-C95 disulfide bridge as predicted by molecular dynamics simulations in this study.

It is important to note that only a few examples exist for the formation of disulfide bridges in soluble protein complexes that reside in the cytoplasm, an observation that has been attributed to the reducing properties of this compartment. For instance, viruses maturing in the cytoplasm can form stable structural disulfide bonds in their coat proteins (Locker and Griffiths, 1999; Hakim and Fass, 2010). Moreover, a number of cytosolic proteins, including phosphatases, kinases and transcriptions factors, have been recognized to be regulated by thiol oxidation and disulfide bond formation, formed as a post-translational modification (Lennicke and Cocheme, 2021). In numerous cases with direct relevance for our studies on FGF2, disulfide bond formation and other forms of thiol oxidation occur in association with membrane surfaces. In fact, many of these processes are linked to the inner plasma membrane leaflet (Nordzieke and Medrano-Fernandez, 2018). Growth factors, hormones and antigen receptors are observed to activate transmembrane NADPH oxidases generating O_2_ ^·-^/H_2_ O_2_ (Brown and Griendling, 2009). For example, the local and transient oxidative inactivation of membrane-associated phosphatases (e.g., PTEN) serves to enhance receptor associated kinase signaling (Netto and Machado, 2022). It is therefore conceivable that similar processes introduce disulfide bridges into FGF2 while assembling into oligomers at the inner plasma membrane leaflet, providing a mechanism that ensures that FGF2 monomers do not prematurely oligomerize in the cytoplasm of cells. In future experiments, it will be important to study disulfide-dependent FGF2 oligomerization in a cellular context, revealing the oxidant and the enzymatic systems that mediate this process at a time scale compatible with the kinetics observed for FGF2 membrane translocation in intact cells (Dimou et al., 2019).

In conclusion, with the current study, important insights were obtained regarding the highly dynamic process of PI(4,5)P_2_-dependent FGF2 oligomerization on membrane surfaces, the key trigger of the central step of the unconventional secretory pathway of FGF2, the remodeling of the plasma membrane lipid bilayer into a lipidic membrane pore with a toroidal architecture. With a number of other cargo proteins secreted by UPS type I pathways in a PI(4,5)P_2_-dependent manner such as HIV-Tat, Tau and homeoproteins (Rayne et al., 2010; Debaisieux et al., 2012; Rabouille, 2017; Katsinelos et al., 2018; Merezhko et al., 2018; Merezhko et al., 2020; Joliot and Prochiantz, 2022; Sparn et al., 2022b), our findings pave the way for a deeper understanding of the general principles of UPS type I pathways of unconventional protein secretion.

## Materials and Methods

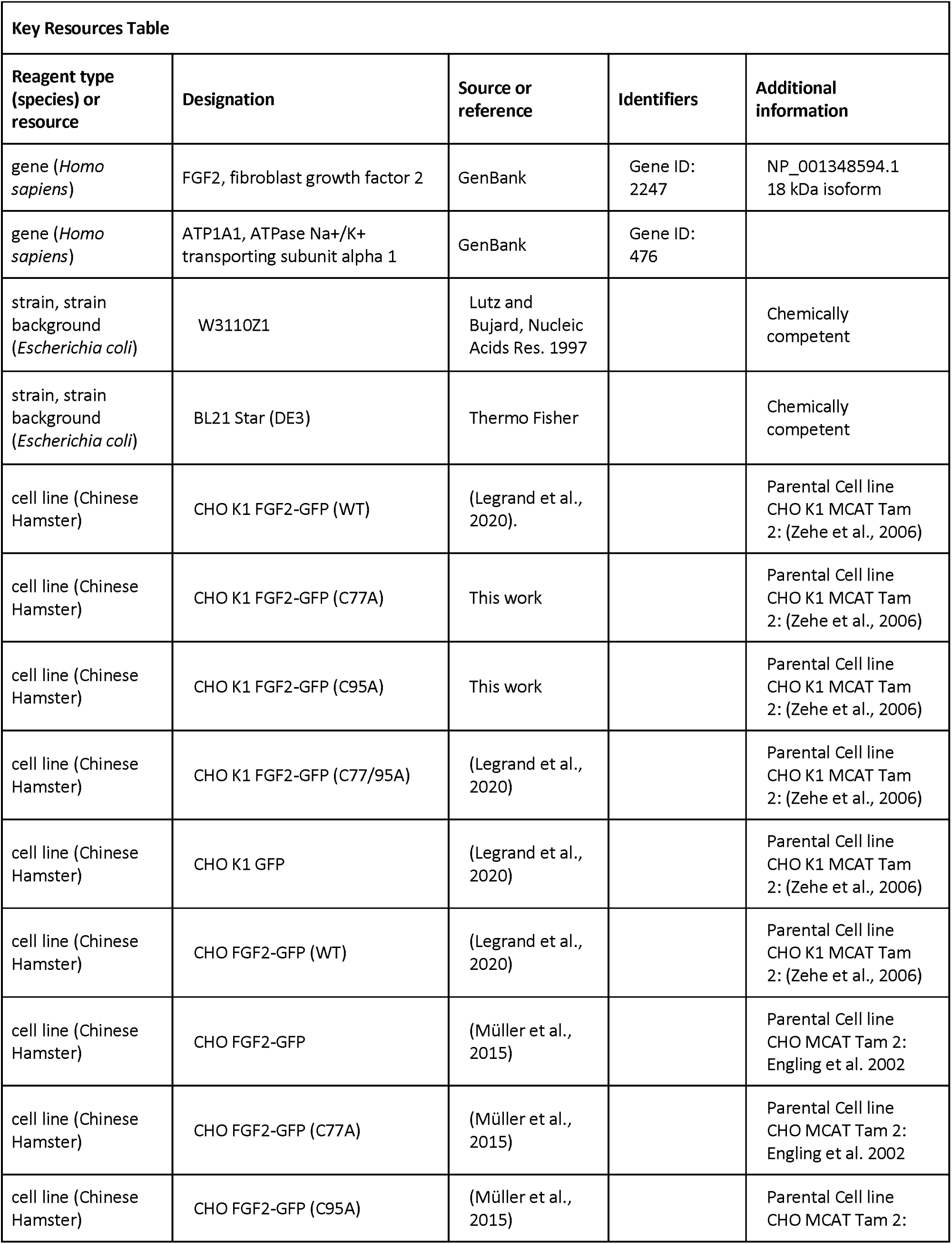
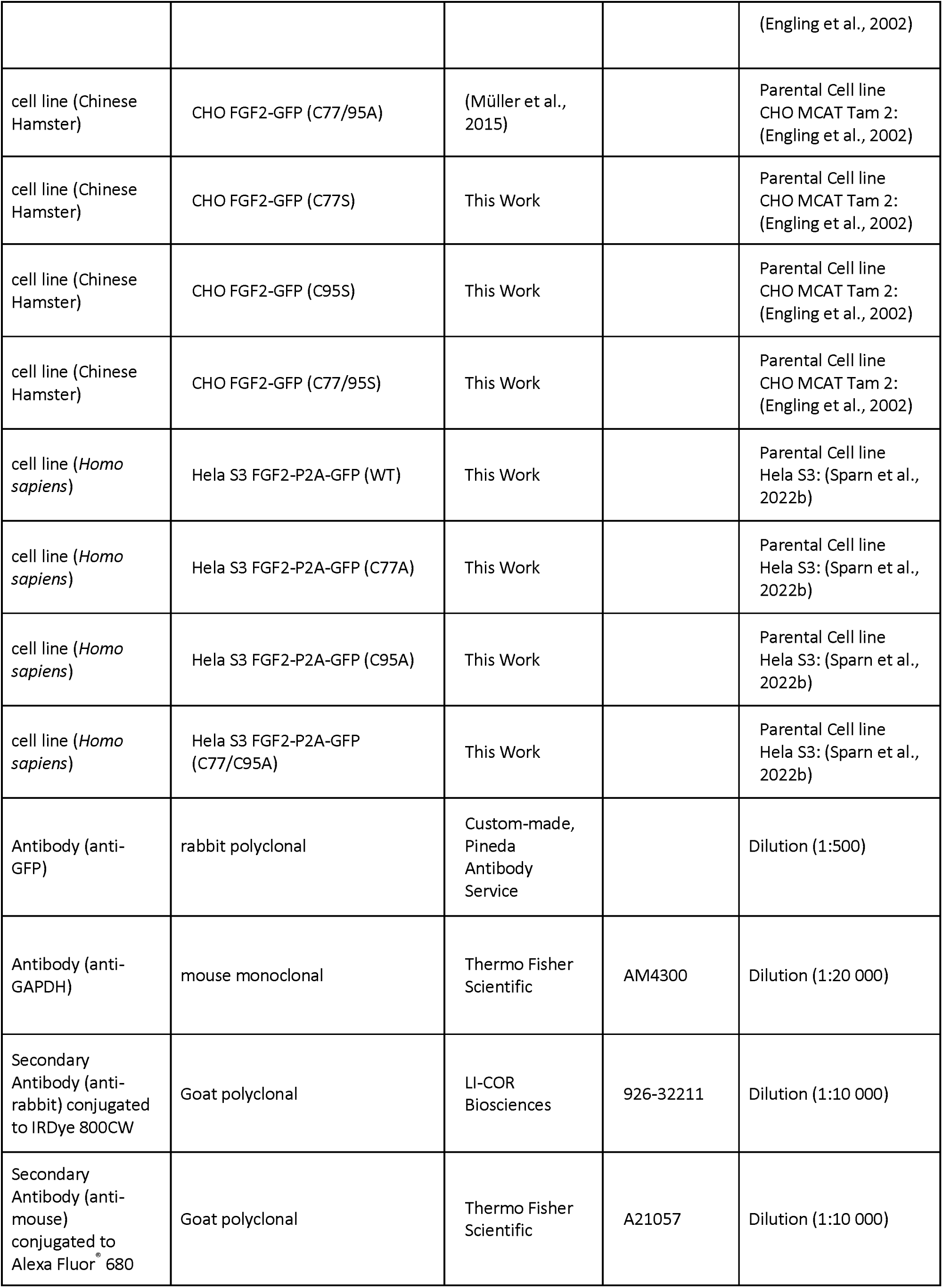
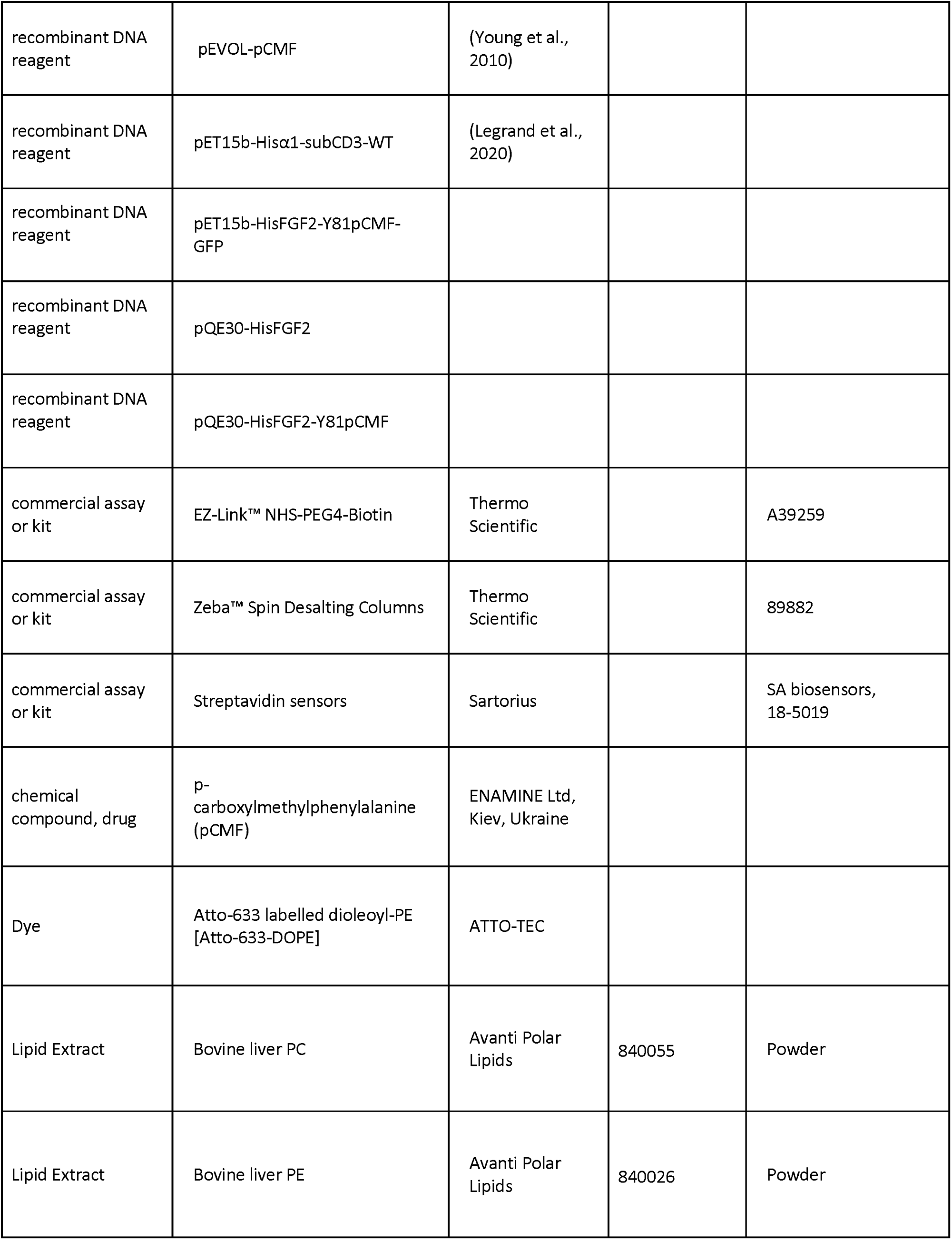
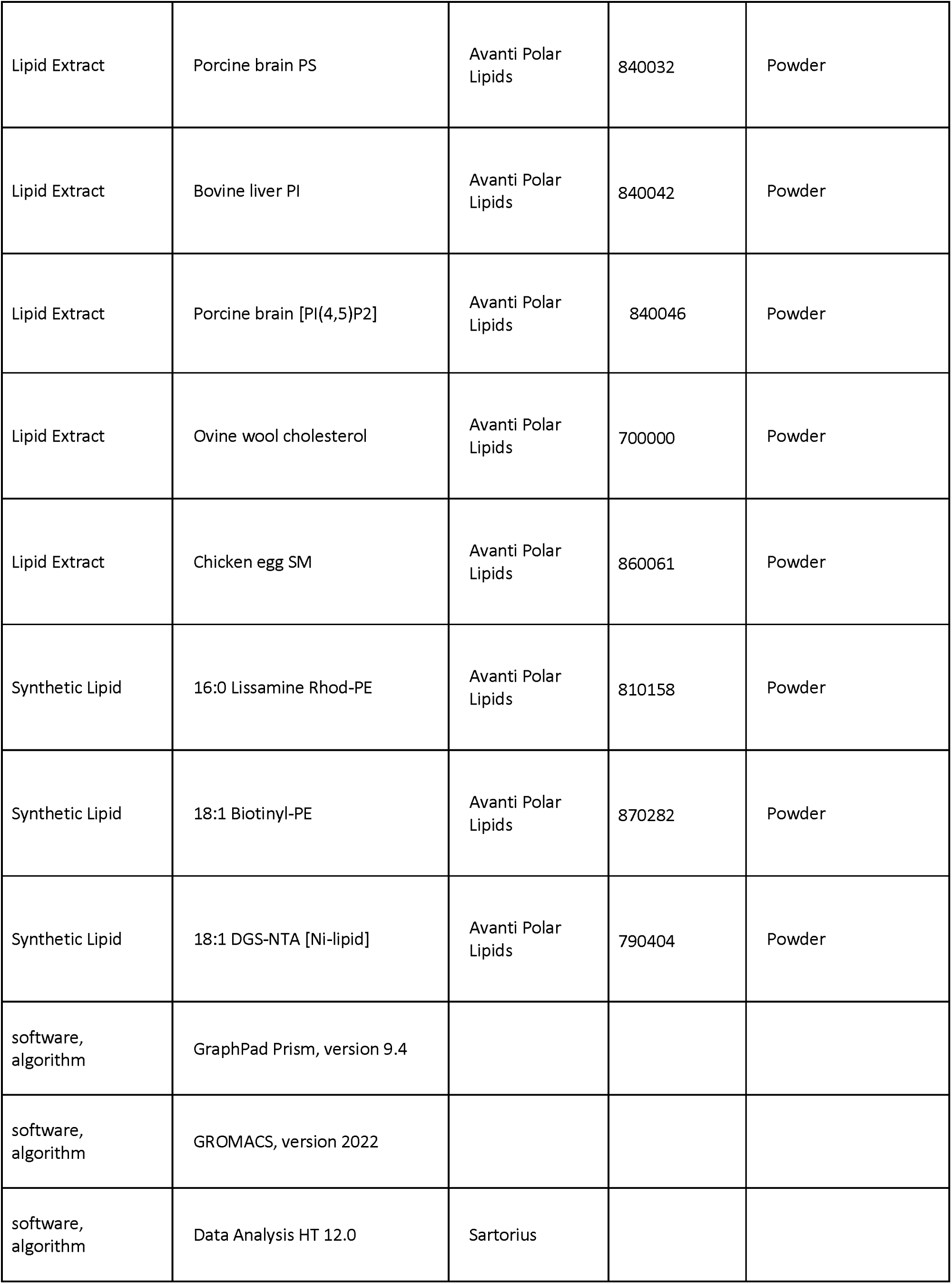
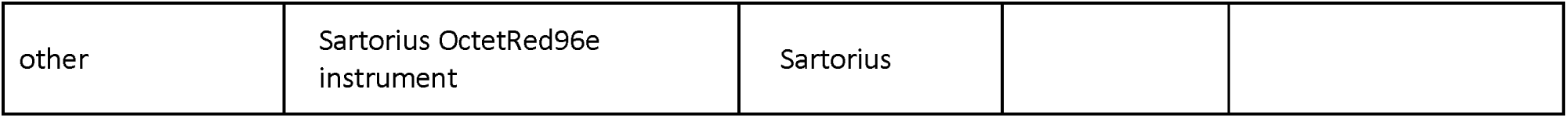

### Cell Culture

CHO, CHO K1 and HeLa S3 cells were cultured in Minimal Essential Medium Eagle (α-MEM) and Dulbecco’s modified Eagle’s medium (DMEM), respectively (Legrand et al., 2020; Lolicato et al., 2022; Sparn et al., 2022a). Both media were supplemented with 10% heat inactivated Fetal Calf Serum (FCS), 100 U/mL penicillin, and 100 μg/mL streptomycin. α-MEM was further supplemented with 2 mM glutamine. Cell lines were grown at 37 °C, in the presence of 5 % CO_2_, and with 95 % humidity. All parental cell lines used in this study were received from the Leibniz Institute DSMZ (German collection of microorganisms and cell cultures GmbH). For human cell lines, their identities were confirmed by STR profiling. CHO K1 cells’s identity and purity were analyzed by a multiplex cell contamination test. All cell lines tested negative for mycoplasma contaminations.

### Generation of stable cell lines

Stable cell lines (CHO and CHO K1) were generated with a retroviral transduction system based on Moloney Murine Leukemia Virus as previously described (Engling et al., 2002). Virus production was performed in HEK293 cells with a stably integrated pVPack-Eco packaging system in its genome as well as the retroviral packaging proteins (EcoPack 2-293 cells). Proteins to be expressed upon induction with doxycycline (like FGF2-GFP), were cloned into the pRevTre2 vector, containing a Tet-response element. Retrovirus production was performed according to the MBS Mammalian Transfection Kit (Agilent Technologies) and virus was harvested after 2 days from confluent cells. CHO and CHO K1 cells constitutively expressing the murine cationic amino acid transporter MCAT-1 (Albritton et al., 1989) and a Tet-On transactivator, rtTA2-M2 (Urlinger et al., 2000), were transduced with the freshly harvested virus. GFP-expressing cells were selected by FACS.

### Gel electrophoresis and Western analyses

Samples were loaded on NuPAGE Bis-Tris 4-12% gradient polyacrylamide gels (Thermo Fisher Scientific). SDS-PAGE was conducted in either MES or MOPS SDS running buffer (Thermo Fisher Scientific) at 200 V for 40 min. As a molecular weight marker, a pre-stained PageRuler was used (Thermo Fisher Scientific). Proteins were then transferred to a methanol-activated PVDF membrane (Millipore) for 1 h at 100 V, in blot buffer (40 mM glycine, 25 mM Tris base, and 20% methanol). After protein transfer, PVDF membranes were blocked for 1 h at room temperature with PBS containing 5% milk. After that, two washing steps of 5 min each were performed in PBS-T (PBS supplemented with 0.05% Tween20) at room temperature prior to the addition of a primary antibody solution (in PBS-T supplemented with 2% BSA and 0.02% Na-azide). After 1 h incubation at room temperature, blots were washed four times with PBS-T prior to the addition of fluorescently-labelled secondary antibodies [goat anti-mouse AlexaFluor 680 (Thermo Fisher Scientific) and goat anti-rabbit IRDye 800CW (LI-COR Biosciences)]. After 30 min incubation at room temperature, blots were washed four times with PBS-T and once with PBS. Imaging was performed with the LI-COR Odyssey^lZI^ CLx system and analyzed with Image Studio Lite (Version 5.0.21, LI-COR Biosciences).

### Cell surface biotinylation

Cell surface biotinylation assays were conducted with stable cell lines producing FGF2 fusion proteins in a doxycyclinbe-dependent manner as described previously (Müller et al., 2015; Legrand et al., 2020; Sparn et al., 2022a). 2 x 10^5^ CHO cells were seeded on 6-well plates 48 h prior the conductance of the assay. These cell lines expressed either wild-type or mutant forms of FGF2-GFP (C77A, C95A, C77/95A) in a doxycycline-dependent manner. Protein expression was induced with the addition of 1 µg/mL doxycycline 24 h after seeding. To perform the biotinylation assay, cells were put on ice and washed twice with PBS supplemented with 1 mM MgCl_2_ and 0.1 mM CaCl_2_. Subsequently, cells were incubated on ice with 1 mg/mL sulfo-NHS-SS-biotin dissolved in incubation buffer (150 mM NaCl, 10 mM triethanolamine, pH 9.0, 2 mM CaCl_2_) for 30 min. Biotinylation was stopped by washing the cells once with quenching buffer (100 mM glycine in PBS supplemented with 1 mM MgCl_2_ and 0.1 mM CaCl_2_) and incubating for an additional 20 min on ice with quenching buffer. Cells were then washed twice with PBS and lysed for 10 min at 37°C in lysis buffer [62.5 mM EDTA pH 8.0, 50 mM Tris-HCl pH 7.5, 0.4% sodium deoxycholate, 1% Nonidet P-40, and protease inhibitor cocktail (Roche)]. Lysed cells were detached by scraping, transferred to fresh 1.5 mL tubes and sonicated in a sonication bath for 3 min. Cell lysates were then incubated for 15 min at room temperature and vortexed every 5 min. Lysates were centrifuged at 18,000 g for 10 min at 4°C to remove cell debris. 5% of the supernatant (15 µL out of 300 µL) was used as a total cell lysate input and boiled at 95°C for 10 min after the addition of SDS sample buffer (40% glycerol, 240 mM Tris-HCl pH 6.8, 8% SDS, 5% β-mercaptoethanol, and bromphenol blue). The remaining 95% of whole cell lysates were incubated using continuous overhead turning at room temperature with Pierce Streptavidin UltraLink Resin (Thermo Fisher Scientific) beads that were previously washed twice with lysis buffer. Washing steps were performed at 3000 x g for 1 min. After 1 h incubation, unbound material was removed via centrifugation (1 min at 3000 x g, RT) and beads were washed once with washing buffer 1 (lysis buffer supplemented with 0.5 M NaCl) and three times with washing buffer 2 (lysis buffer containing 0.1% NP-40, supplemented with 0.5 NaCl). Bound material was then eluted via boiling at 95°C for 10 min with SDS sample buffer. Input and eluate samples were then separated via SDS-PAGE and analyzed by Western blot analysis. Western analysis was conducted for FGF2-GFP [detected with an anti-GFP polyclonal rabbit antibody (Custom made, Pineda Antibody Service)] and GAPDH [detected with a mouse monoclonal antibody (Thermo Fisher Scientific)].

### Real-time Single Molecule TIRF analyses in cells

Real-time single molecule TIRF assays to quantify FGF2 recruitment at the inner plasma membrane leaflet under various experimental conditions was conducted as described previously (Dimou et al., 2019; Legrand et al., 2020; Lolicato et al., 2022). CHO K1 expressing FGF2 variants (wild-type, C77A, C95A, or C77/95A) as GFP fusion proteins or GFP alone in a doxycycline-dependent manner were cultivated on 8-well glass bottom µ-slides (ibidi) for 24 hours before experiments were started. FGF2-GFP expression levels were kept low in the absence of doxycycline to allow for single molecule measurements. Before imaging, cells were washed twice with Live Cell Imaging Solution (Thermo Fisher Scientific). Both wide-field and TIRF imaging were performed using an inverted Olympus IX81 xCellence TIRF microscope equipped with an Olympus PLAPO 100x 1.45 NA Oil DIC objective lens and a Hamamatsu ImagEM Enhanced (C9100-13) camera. Wide-field images were exploited to detect the frames of each cell, and GFP fluorescence was excited with an MT 20 illumination system. TIRF time-lapse videos were analyzed to detect single FGF2-GFP particles recruited at the inner plasma membrane leaflet, and GFP fluorescence was excited with an Olympus 488 nm, 100 mW diode laser. Both wide-field and TIRF images were recorded with the Olympus xCellence software, saved as Tagged Image File Format (TIFF), and analyzed via Fiji (Schindelin et al., 2012). FGF2-GFP recruitment efficiencies were quantified after normalizing for both FGF2-GFP expression levels (quantified for each analyzed cell at the first frame of TIRF time-lapse videos) and surface area (µm^2^). To quantify single FGF2-GFP particles recruited at the inner plasma membrane leaflet, the Fiji plugin TrackMate was employed (Tinevez et al., 2017). For every representative image shown, background fluorescence was subtracted.

### Transient expression of FGF2 variant forms and cross-linking experiments in in HeLa S3 cells

HeLa S3 cells were seeded on 6-well plates and transfected after 24 hours of incubation with a plasmid based on pcDNA 3.1 encoding a FGF2-P2A-GFP fusion protein, using the FuGENE transfection reagent (Promega). Following 48 hours of further incubation, whole cell lysates were prepared in a detergent-containing buffer [50 mM HEPES (pH 7.4), 1% Nonidet P-40, 0.25% sodium deoxycholate, 50 mM NaCl, 10% Glycerol, Halt^TM^ Protease and Phosphatase Inhibitor Cocktail (ThermoFisher)]. Lysates were incubated on ice for 30 min followed by centrifugation at 20,000 x g for 10 min at 4 °C. Supernatants were subjected to crosslinking reactions using (i) BMH (bismaleimidohexane), (ii) PMPI (N-p-maleimidophenylisocyanate), or (iii) BMOE (bismaleimidoethane) that were prepared in DMSO and further diluted to a final concentration of 0.2 mM. Following incubation for 30 min at room temperature in the dark, excess amounts of crosslinkers were quenched for 15 min with 50 mM DTT. FGF2 crosslinking products were separated by SDS-PAGE and quantified by Western analysis using anti-FGF2 polyclonal rabbit antibody (Custom made, Pineda Antibody Service) and goat anti-rabbit IRDye 800CW (LI-COR Biosciences) as secondary antibody (Engling et al., 2002; Backhaus et al., 2004; Schäfer et al., 2004). The ratio of monomeric versus dimeric species of FGF2 was quantified using a LI-COR Odyssey imaging system. The corresponding data were statistically evaluated as explained in the legend to Fig. 3.

### Recombinant Proteins

His-tagged variants of FGF2 [WT, C77A, C95A, C77/95A, K54/60A, K54/60A-C77A (Biolayer Interferometry)], FGF2-Y81pCMF (WT, C77A, C95A, C77/95A (Membrane Pore Formation Assay)] (both pQE30), FGF2-Y81pCMF-GFP [(WT, C77A, C95A, C77/95A (Dual-color FCS Measurement and Translocation Assay)], as well as Hisα1-subCD3-WT (Biolayer Interferometry) (both pET15b) were expressed in Escherichia coli strains W3110Z1 or BL21 Star (DE3), respectively. For incorporation of the unnatural amino acid p-carboxylmethylphenylalanine (pCMF; custom synthesis by ENAMINE Ltd., Kiev, Ukraine), codon 81 (tyrosine) was replaced by an amber stop codon. Transformation of a strain carrying the pEVOL-pCMF plasmid resulted in expression of recombinant FGF2-Y81pCMF (Young et al., 2010). Recombinant proteins were purified to homogeneity in three steps using Ni-NTA affinity chromatography, heparin chromatography (except ATP1A1subCD3), and size exclusion chromatography using a Superdex 75 column (Steringer et al., 2017). Protein purity was determined by sodium dodecyl-sulfate polyacrylamide gel electrophoresis (SDS-PAGE) under reducing conditions. For each protein, 3 μg were loaded. Protein patterns were analyzed by Instant Blue© Coomassie staining (abcam).

### Lipids

Membrane lipids from natural extracts (bovine liver phosphatidylcholine [PC], bovine liver phosphatidylethanolamine [PE], porcine brain phosphatidylserine [PS], bovine liver phosphatidylinositol [PI], porcine brain phosphatidylinositol-4,5-bisphosphate [PI(4,5)P_2_], ovine wool cholesterol, and chicken egg sphingomyelin [SM]) as well as synthetic products (16:0 Lissamine Rhodamine-labeled PE [Rhod.-PE], 18:1 Biotinyl-PE [Biotinyl-PE], and 18:1 DGS-NTA (Ni) [Ni-lipid]) were purchased from Avanti Polar Lipids. In addition, Atto-633 labelled dioleoyl-PE [Atto-633-DOPE] was purchased from ATTO-TEC.

### Preparation of giant unilamellar vesicles (GUVs)

GUVs with a plasma membrane like lipid composition consisting of 30 mol% cholesterol (Chol), 15 mol% sphingomyelin (SM), 34 mol% phosphatidylcholine (PC), 10 mol% phosphatidylethanolamine (PE), 5 mol% phosphatidylserine (PS), 5 mol% phosphatidylinositol (PI) and 1 mol% Biotinyl-PE (Avanti Polar Lipids) were generated based on electro-swelling using platinum electrodes (García-Saéz et al., 2009). GUVs were supplemented with either phosphatidylinositol-4,5-bisphosphate PI(4,5)P_2_ or a Ni-NTA lipid at 2 mol % at the expense of PC as indicated. For visualization either 0.05 mol% rhodamine B-labelled PE for FGF2 translocation assays or 0.002 mol% Atto-633 labelled dioleolyl-PE (Atto-633-DOPE, ATTO-TEC) for Dual-color FCS measurements was added. The dried lipid film was hydrated with a 300 mM sucrose solution (300 mOsm/kg, Wescor Vapro). Where indicated, long-chain heparins (50 mM; based on disaccharide units) were included in the lumen of GUVs in order to mimic heparan sulfates. Swelling was conducted at 45°C [10 Hz, 1.5 V for 50 min (without heparin) or 70 min (with heparin), 2 Hz, 1.5 V for 25 min]. In order to remove excess amounts of heparin and sucrose, GUVs were gently washed twice with buffer B (25 mM HEPES pH7.4, 150 mM NaCl, 310 mOsmol/kg) and collected via centrifugation (1200 x g; 25°C; 5 min). Imaging chambers (LabTek for FGF2 translocation assays, ibidi for Dual-color FCS) were incubated sequentially with 0.1 mg/ml Biotin-BSA (Sigma A8549) and 0.1 mg/ml Neutravidin (Thermo Fisher Scientific A2666) in buffer B.

### Dual-color FCS measurement

The oligomeric state of His-tagged FGF2-Y81pCMF-GFP variants (WT, C77A, C95A and C77/95A) was determined by z-scan FCS measurements performed on a home-built confocal microscope system consisting of an Olympus IX71 inverted microscope body (Olympus, Hamburg, Germany) with a 3D piezo positioner from Physik Instrumente (P-562.3CD stage controlled via E-710.3CD controller) and pulsed diode laser heads PicoTA-532, LDH-P-C-470 and LDH-P-635 controlled via PDL 828 Sepia II laser driver (all devices from PicoQuant, Berlin, Germany) at room temperature. The lasers were pulsing alternately in order to avoid artefacts caused by signal bleed-through. The beam was coupled to a single polarization maintaining single mode fibre, collimated by an air space objective (UPlanSApo 4x, NA 0.16) and directed towards the water immersion objective (UPlanSApo 60x w, NA 1.2) by a Chroma ZT375/473/532/635rpc quad-band dichroic mirror. The collected signal, which passed through a 50 µm diameter hole in the focal plane, was split between two SPAD detectors ($PD-50-CTC, MicroPhotonDevices, Bolzano, Italy) using T635lpxr splitter and HQ515/50 (FGF2-GFP) and HQ697/58 (DOPE-Atto 633) filters mounted in front of each detector. Time-tagged time-resolved single photon counting data acquisition was performed by HydraHarp400 Multichannel Picosecond Event Timer & TCSPC Module controlled via SymPhoTime software (both from PicoQuant, Berlin, Germany). The laser intensity at the back aperture of the objective was kept below 10 mW for each laser line. The z-scan was performed on the top of a selected GUV. The GUV was positioned into the laser beam waist of 470 and 635 nm lasers and moved 1.5 μm below the waist and consequently vertically scanned in 10 to 15 steps (150 nm spaced). At every position, a 60-second-long dual-color FCS measurement was performed.

### Determination of the FGF2 oligomeric state on a single GUV

The oligomeric state of variant forms of FGF2-Y81pCMF-GFP (WT, C77A, C95A and C77/95A) was assessed by comparing the brightness of the protein oligomer with the brightness of a defined monomer. The auto-correlation curves obtained from FCS measurements were fitted by a model assuming two-dimensional diffusion in the membrane (bound FGF2-GFP and DOPE-Atto-633), free three-dimensional diffusion in the solution (FGF2-GFP in the bulk) and transition of the dye to the triplet state (Widengren et al., 1995):

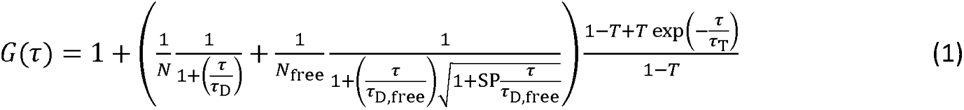

The τ represents the lag-time, N and N_free_ are the number of membrane-bound and free dye molecules in the confocal volume, τ_D_ and τ_D,free_ the diffusion times of membrane bound and membrane free dye, SP the structure parameter, *T* the fraction of the dye in the triplet state and τ_T_ the lifetime of the triplet state. The auto-correlation curves were recorded for both DOPE-Atto633, reporting on the quality of the membrane, and FGF2-GFP that were analyzed to determine the brightness of an FGF2 oligomer. In the beam center, the fluorescent signal coming from the solution is negligible. This simplifies the above equation into:

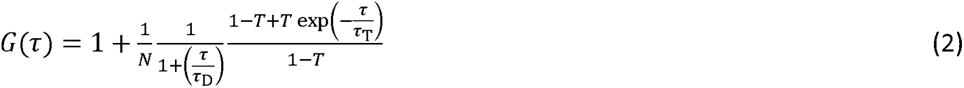

Obtaining reliable output parameters requires a precise focus into the beam center, which is achieved by successive vertical scanning of the membrane along the z axis. For further analysis, the position of the membrane with the minimum in N and the corresponding average intensity in counts per second ⟨*I*⟩, was used (Benda et al., 2003). The average oligomeric state of FGF2 on individual GUV was calculated by comparing the brightness of an oligomer *ɸ*(oligo) to that of a monomer *ɸ*(mono). The brightness of an oligomer was calculated as 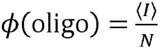. The brightness of a monomer *ɸ*(mono) is obtained in a similar way, however, the presence of one labelled molecule of FGF2-Y81pCM-GFP in a cluster must be ensured. To fulfil this requirement, His-FGF2-Y81pCMF-C77/95A-GFP was diluted by its unlabeled variant (His-FGF2-Y81pCMF-C77/95A) at a ratio of 1:10. Alternatively, recombinant His-FGF2-Y81pCMF-C77/95A-GFP which binds to DGS-NTA containing lipid bilayers as a dimer at maximum was used (Müller et al., 2015; Steringer et al., 2017; Sachl et al., 2020). Finally, the average oligomeric state was calculated as

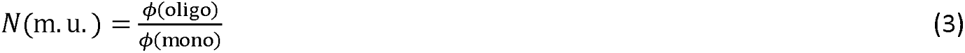

### Imaging and quantification of FGF2 membrane translocation using GUVs

For FGF2 membrane translocation assays, GUVs were incubated for 3 h with a small fluorescent tracer (Alexa647) and His-tagged FGF2-Y81pCMF-GFP variants (WT, C77A, C95A, C77/95A) at a final concentration of 200 nM in buffer B (25 mM HEPES pH 7.4, 150 mM NaCl, 310 mOsmol/kg) as described previously (Steringer et al., 2017). Confocal images were recorded at room temperature in multitrack mode using Zeiss LSM510 confocal fluorescence microscopes (Carl Zeiss AG, Oberkochen, Germany) using a plan apochromat 63×, NA 1.4 oil immersion objective. Pinholes of the tracks were optimized to 1.2 mm. In order to measure (i) GFP-, (ii) Rhodamine-PE-, and (iii) Alexa647-derived signals, samples were excited with (i) an argon laser (488 nm), (ii) a He-Ne-laser (561 nm), or (iii) a He-Ne laser (633 nm). The emission signal was detected after (i) a band pass (BP) filter (505–530 nm), (ii) a BP filter (560–615 nm), or (iii) a long pass (LP) filter (>650 nm). Images were recorded in 8-bit grayscale. The luminal fluorescence of individual GUVs was measured and normalized to the fluorescence intensity of the surrounding buffer. For each experimental condition, 20 to 120 individual GUVs were analyzed using ImageJ software (reference) (http://rsbweb.nih.gov/ij/). To allow for statistical analysis of membrane pore formation and FGF2-GFP membrane translocation across the population of GUVs, thresholds were defined to classify individual GUVs. When the inside-to-outside fluorescence ratio of the Alexa647 tracer was ≥0.6, GUVs were classified as vesicles containing membrane pores. Similarly, when the inside-to-outside ratio of GFP fluorescence was ≥1.6, the corresponding GUVs were classified as vesicles where FGF2-GFP membrane translocation into the lumen had occurred. Statistical analyses are based on two-tailed, unpaired t-test using Graph Pad Prism, Version 9.4.1.

### Preparation of large unilamellar vesicles and membrane pore formation assays

Liposomes with a plasma membrane-like lipid composition consisting of 50 mol% Chol, 12.5 mol% sphingomyelin (SM), 15.5 mol% PC, 9 mol% PE, 5 mol% PS, 5 mol% PI, 2 mol % of PI(4,5)P_2_, and 1 mol% Rhod.-PE were prepared as described previously (Temmerman et al., 2008; Temmerman and Nickel, 2009; Steringer et al., 2012; Müller et al., 2015). In brief, chloroform-dissolved lipid mixtures were first dried under a gentle nitrogen stream and further dried under vacuum for 1.5 h to yield a homogeneous lipid film. Lipids were resuspended in buffer A [100 mM KCl, 25 mM HEPES, pH 7.4, 10% (w/v) sucrose] supplemented with 100 µM concentration of the membrane-impermeant fluorophore 5(6)-carboxyfluorescein (Sigma) at 45 °C to form liposomes with a final lipid concentration of 8 mM. Liposomes were subjected to 10 freeze/thaw cycles (50°C/liquid nitrogen) and to 21 size-extrusion steps (400 nm pore size; Avanti Polar Lipids mini-extruder). Liposome preparations were analyzed by dynamic light scattering (Wyatt), indicating a range of 200 to 400 nm in diameter. To remove extraluminal 5(6)-carboxyfluorescein, liposomes were diluted in buffer B (150 mM KCl, 25 mM HEPES, pH 7.4) and collected by centrifugation at 15,000 g for 10 min at 20 °C followed by size exclusion chromatography using a PD10 column (GE Healthcare). Importantly, this column was operated in buffer C [150 mM KCl, 25 mM HEPES, pH 7.4, 10% (w/v) sucrose, 2% (w/v) glucose] that was titrated with glucose to reach iso-osmolality (840 mOsm/kg, Wescor Vapro). After incubation with various forms of His-tagged FGF2 (WT, C77A, C95A, C77/95A) at a final concentration of 2 µM, fluorescence dequenching was measured using a SpectraMax M5 fluorescence plate reader (Molecular Devices). At the end of each experiment, Triton X-100 [0.2% (w/v) final concentration] was added to measure maximal dequenching used to normalize data.

### Biolayer interferometry (BLI) to quantify α1 interaction with FGF2 variants

Measurements were conducted on a Sartorius OctetRed96e instrument using Streptavidin sensors (SA, Sartorius 18-5019). This type of sensor was chosen because the nonspecific binding control showed less than 10% of the corresponding signal, which is a prerequisite of BLI measurements. Hisα1-subCD3-WT (Legrand et al., 2020) was biotinylated with EZ-Link™ NHS-PEG4-Biotin (Thermo Scientific™ A39259) in a 1:1.5 ratio at 25°C for 30 min. Free Biotin was removed using Zeba™ Spin Desalting Columns (Thermo Scientific™ 89882) according to the manufacturer’s instruction. In order to find the optimal ligand concentration, a loading scout was performed. In the following experiments 4 µg/ml (150 nM) biotinylated Hisα1-subCD3-WT was loaded for 20 min. Next, FGF2-WT was titrated (1:2 dilution series starting at 1000 nM) in order to determine K_D_, k_a_ and k_d_ values. For comparison of FGF2 variants, a concentration of 1000 nM was selected to make use of the full dynamic range of the assay. Kinetic assay step times were as follows: Equilibration (5 min), Loading of ligand (20 min), Wash (5 min), Baseline (3 min), Association (10 min), Dissociation (10 min). All steps were performed in assay buffer [0.02 %(w/v) Tween-20, 0.1 %(w/v) BSA in PBS] at 25°C while shaking (1000 rpm).

Data were analyzed with Data Analysis HT 12.0 software version 12.0.2.59 (Sartorius). In brief, a single referencing setup was used (drift) where a loaded sensor was run in parallel in an assay buffer containing a reference well. Later the resulting signals were subtracted from all sample well data. According to kinetic measurement guidelines data curves were aligned in Y using the function “Average Baseline Step”, inter-step correction “Dissociation Step”, and Savitzky-Golay-Filter. K_D_, k_a_ and k_d_ values were calculated for FGF2-WT to Hisα1-subCD3-WT by globally fitting Association and Dissociation using a 1:1 model. Curves that showed a high residual were excluded according to the manufacturer’s guidelines. Mean with standard deviations of K_D_, k_a_, and k_d_ values (n = 3) are given.

### 360 degree analysis: Sampling of the dimerization interface through atomistic molecular dynamics simulations

To obtain a coverage of possible FGF2-FGF2 dimerization interfaces compatible with a membrane-bound state, we performed 360 atomistic molecular dynamics (MD) simulations. The goal was to find out all dimerization interfaces where C95 is involved. Both FGF2 monomers were placed on the surface of a model membrane composed of POPC enriched with 30 mol% cholesterol. Initially, both monomers pointed in the same direction, based on our previous work where we observed a high-affinity orientation for FGF2 characterized by strong PI(4,5)P_2_-mediated binding to the membrane surface (Steringer et al., 2017, eLife). In this orientation, the known PI(4,5)P_2_ binding site of FGF2 (K127, R128, K133; Temmerman et al., 2008, Traffic) is directly against the membrane surface. We then started with a situation where the C95 residue of the first FGF2 monomer interacted through contact with the surface of the second FGF2 monomer. Next, with the first monomer held in place with its C95 pointed directly toward the second monomer, this second FGF2 monomer was rotated one degree at a time around the z-axis (membrane normal direction), thanks to which we obtained 360 structures in which the FGF2 monomers were in different orientations with respect to each other, and we were able to create initial structures to elucidate all dimerization interfaces where C95 is involved. In each of these cases, the monomers were placed 0.8 nm apart so that in the chosen dimer structure they hit the van der Waals and Coulomb cutoff of the force field (1.2 nm), which ensured the proximity of the monomers but without a strongly bound initial state. For each of the 360 cases, a 500 ns simulation was performed with the previously defined settings. The simulation data was analyzed using machine learning tools. Finally, it is worth noting that checking all possible dimerization interfaces was not the goal of this work, because its implementation with the same resolution would not have been feasible – it would have required 360 x 360 simulations. Despite this, due to the rotation and diffusion of the protein monomers during the simulations, these simulations also sampled to some extent those dimerization interfaces where C95 was not involved.

### Machine learning based analysis

To identify the prevalent dimerization interface in the 360-degree simulations described above, we used the Bayesian Gaussian Mixture Model (GMM) based clustering in the structural space spanned by the simulations. The simulation structures were rotationally and translationally fitted to one of the two monomers and its associated PI(4,5)P_2_ lipid to orient the other monomer in a fixed reference frame. For practical reasons, the dimensionality of the structural space from the fitted trajectory was first reduced using an Artificial Neural Network (ANN) based auto-encoder (AE) as this workflow allows for more robust clustering. This fitted trajectory was then divided into two equal parts, the training set and the cross-validation set. The training set was used to construct the model weights. The reconstruction error for the cross-validation set in combination with an orthogonal loss was used to train the model over 50 epochs with early stoppings to prevent overfitting (Wang et al., 2019). The orthogonal loss function was added to reduce the correlation in the encoded layer neurons. The input layer for the AE was supplied by the coordinates of the C-alpha atoms of the two monomers. The autoencoder was constructed using a total of 5 dense hidden layers. The two encoding layers were constructed with 1024 neurons followed by the encoded layer with two neurons. These three layers formed in effect the Encoder part of the architecture. The two decoding layers were constructed with 1024 neurons each. The ReLu activation function was used for all neurons. Dropout regularization was used for each layer to enhance the sparsity of the dimensionality reduction model, and L2 regularization was added to avoid over-reliance on highly activating neurons. The pytorch library was used to construct the AE (Paszke et al., 2019).

Clusters were identified using Bayesian-GMM in the 2D space encoded by the AE. A ‘full’ type covariance matrix was used to identify the parameters of the Gaussian distributions used to create the clusters. The initial guesses for the expectation-maximization algorithm for the GMM were placed using the K-means algorithm. Through visual testing, an 8-component model was finally chosen as a robust representation of the clustering in the encoded space. The scikit-learn package was used to train the GMMs (Pedregosa et al., 2011).

### In silico protein-protein docking studies

We conducted in silico protein-protein docking studies using the ROSETTA 2018 package (Gray et al., 2003; Wang et al., 2005; Wang et al., 2007; Chaudhury and Gray, 2008). Our starting structure was the truncated FGF2 monomer [PDB id: 1BFF; (Kastrup et al., 1997)] from residue 26 to 154 without the flexible N-terminus. We positioned two monomers so that the C95 residues faced each other, with the experimentally known PI(4,5)P_2_ binding pocket residues oriented towards the same side to allow for simultaneous membrane interactions. To generate diversity, we randomly perturbed one FGF2 monomer by 3 Å translation and 8° rotation before the start of each docking simulation, resulting in 500 structures. We ranked these structures based on the interface score. The top 5 % ranked structures were clustered based on the RMSD value for the FGF2 dimer using the Gromos algorithm (Daura et al., 1999). Two structures were considered neighbors if their RMSD value was within 0.6 nm. Finally, we selected the most representative structure from the four-ranked clusters and simulated it in atomistic molecular dynamics simulations in water as a covalently linked disulfide-bridged dimer.

### Atomistic molecular dynamics simulations

All simulations were carried out with atomic level models. These molecular dynamics (MD) simulations were performed using the CHARMM36m force field for lipids and proteins, the CHARMM TIP3P force field for water, and the standard CHARMM36 force field for ions (Huang et al., 2017). All simulations were carried out using the GROMACS 2022 simulation package (Abraham et al., 2015). For FGF2, we used the crystal structure of residues 26–154 of the monomeric form of FGF2 [PDB id: 1BFF; (Kastrup et al., 1997)], with the N- and C-termini modeled as charged residues. All systems were first energy-minimized in a vacuum and then hydrated as well as neutralized by adding an appropriate number of counter ions and 150 mM potassium chloride to mimic experimental conditions. Next, an equilibration step was performed to keep the temperature, pressure, and number of particles constant (NpT ensemble). During this step, proteins were restrained in all dimensions, while the first heavy atom of each lipid (if present) was restrained in the xy-plane of the membrane with a force constant of 1,000 kJ·mol^-1^·nm^-2^. The Nose-Hoover thermostat maintained the temperature at 310 K with a time constant of 1.0 ps (Evans and Holian, 1985). The pressure of 1 atm was kept constant using the Parrinello–Rahman barostat with a time constant set to 5.0 ps and an isothermal compressibility value of 4.5 × 10^−5^ bar^−1^ (Parrinello and Rahman, 1981). The isotropic pressure-coupling scheme was used for protein-only simulations, while the semi-isotropic scheme was used in the presence of a membrane. We used the Verlet scheme for neighbor searching with an update frequency of once every 20 steps (Verlet, 1967). Electrostatic interactions were calculated using the Particle Mesh Ewald method (Darden et al., 1993) with 0.12 nm spacing, a tolerance of 10^-5^ and a cut-off of 1.2 nm. Periodic boundary conditions were applied in all directions. The simulations were carried out using a time step of 2 fs until 1000 ns were reached.

To generate seven C95-C95 dependent FGF2-FGF2 interfaces (protein-only MD simulations), we used three different techniques. First, we extracted one interface from a previous MD simulation snapshot that featured three monomers bound to PI(4,5)P_2_ on the membrane surface (Steringer et al., 2017). Next, we employed ROSETTA software to generate four more initial structures (with the above protocol) from local protein-protein docking simulations. Finally, we obtained the last two interfaces from an AlphaFold2-multimer v3 prediction using ColabFold v1.5.2 default parameters (Mirdita et al., 2022). The two C95 residues were then covalently linked to generate the structure and topology files of the seven disulfide bridge dimers, which were then subjected to 1-microsecond-long MD simulations in water using the CHARMM-GUI web interface (Lee et al., 2016). The final structure of one of the previous simulations, which showed a stable dimer in the presence of two ion pairs (E86 – K118 and E99 – K88), was randomly placed in ten different orientations 2 nm away from a POPC:cholesterol (70:30) membrane surface containing 2 PI(4,5)P_2_ molecules (protein-membrane MD simulations). These configurations were then simulated for 1 microsecond.

### Free energy calculations

We employed an atomistic resolution and used the umbrella sampling method (Torrie and JP, 1974; Torrie and Valleau, 1977) to calculate the potential of mean force (PMF) for the binding of a disulfide-bridged FGF2 dimer on a model membrane surface. The initial configuration for each of the 45 umbrella windows was taken from unbiased MD simulations. The reaction coordinate was set as the center of mass distance between the FGF2 dimer and the phosphate atoms of one leaflet. We simulated each window for 120 ns with a harmonic restraint force constant of 2000 kJ·mol^-1^·nm^-2^, spaced 0.1 nm apart. The first 50 ns of the simulations were discarded as the equilibration phase. The Weighted Histogram Analysis Method was used to reconstruct the free energy profiles (Meng and Roux, 2015). The statistical error was determined with 200 bootstrap analyses (Rubin, 1981).

### Chemical cross-linking and peptide purification

For cross-linking, 0.5 mM of freshly resuspended Disuccinimidyldibutyric urea (DSBU, Thermo Fisher Scientific, 50 mM stock solution in anhydrous DMSO) was added to 0.5 mg/ml of recombinant His-tagged FGF2 (HEPES-Buffer: 10 mM HEPES, 50 mM NaCl, 1 mM DTT, pH 7.5, 70 µl, pH 7.4 – 8.2) with and without PI(4,5)P_2_-liposomes and incubated at room temperature. After 20 min the reaction was quenched by the addition of tris buffer (1 M, pH 8.0) to a final concentration of 20 mM at room temperature for 30 min. The reaction volume was then doubled by the addition of 100 mM ammonium bicarbonate (ABC) buffer and both, crystalline urea and dithiothreitol (DTT) were added to a final concentration 8 M and 5 mM respectively to denature and reduce the cross-linked proteins at 56 °C for 30 min. Subsequently, the samples were alkylated by the addition of 8 mM iodoacetamide for 30 min in the dark at room temperature. To start the digestion of the cross-linked proteins, Lys-C (0.25 µg/µl, 2µl) was added for 4 h at 37 °C. Afterwards, the reaction solution was diluted to a final concentration of 2 M Urea by the addition 100 mM ABC and the samples were then further digested by the addition of trypsin (1 µg/µl, 2 µl) at 37 °C over night. After digestion, the reaction was stopped by the addition of formic acid to a final concentration of 1 % and the resulting peptide mixtures were desalted and purified by in-house made C18-StageTips (Rappsilber et al., 2007), eluted and dried under vacuum.

### Liquid chromatography and cross-linking mass spectrometry data analysis

Dried peptides were reconstituted in 10 µl of 0.05% trifluoroacetic acid (TFA), 4% acetonitrile, and 5 µl were analyzed by a Ultimate 3000 reversed-phase capillary nano liquid chromatography (LC) system connected to a Q Exactive HF mass spectrometer (Thermo Fisher Scientific). Samples were injected and concentrated on a trap column (PepMap100 C18, 3 µm, 100 Å, 75 µm i.d. x 2 cm, Thermo Fisher Scientific) equilibrated with 0.05% TFA in water. After switching the trap column inline, LC separations were performed on a capillary column (Acclaim PepMap100 C18, 2 µm, 100 Å, 75 µm i.d. x 50 cm, Thermo Fisher Scientific) at an eluent flow rate of 300 nl/min. Mobile phase A contained 0.1 % formic acid in water, and mobile phase B contained 0.1% formic acid in 80 % acetonitrile / 20% water. The column was pre-equilibrated with 5% mobile phase B followed by an increase of 5–44% mobile phase B in 130 min. Mass spectra were acquired in a data-dependent mode with a single mass spectrometer (MS) survey scan (375–1575 m/z) with a resolution of 120,000 and MS/MS scans of the 10 most intense precursor ions both with a resolution of 60,000. The dynamic exclusion time was set to 30 seconds and automatic gain control was set to 3×10^6^ and 1×10^5^ for MS and MS/MS scans, respectively. Fragmentation was induced by HCD with a stepped collision energy (21, 27, 33) and MS/MS spectra were recorded using a fixed first mass of 150 m/z. The acquisition of MS/MS spectra was only triggered for peptides with charge states 4+ to 7+.

The acquired RAW files were converted into Peak lists (.mgf format) using Proteome Discoverer (Thermo, version 2.1) containing CID-MS2 data. The CID-MS2 spectra were deconvoluted with the add-on node MS2-Spectrum Processor using default settings. For the main search of the cross-linked peptides the in-house developed algorithm XlinkX v2.0 was used (Liu et al., 2017). The following search parameters were used: MS1 precursor ion mass tolerance: 10 ppm; MS2 fragment ion mass tolerance: 20 ppm.; fixed modifications: Cys carbamidomethylation; variable modification: Met oxidation; enzymatic digestion: Trypsin; allowed missed number of missed cleavages, 3. All MS2 spectra were searched against a concatenated target-decoy databases generated based on the wt and mutant sequences of the FGF2 protein. Cross-links were reported at a 2% FDR based on a target decoy calculation strategy (Liu et al., 2017). The detected cross-links were mapped on crystal structure PDB (5×10) by setting the allowed distance constraint to the typical range of the used cross-linker (DSBU = ∼25 Å).

### Structural modeling of FGF2-Halo dimers into subtomogram average electron density maps

A FGF2-HALO dimer structure was generated that fits well into the 3D density using a structural template of AlphaFold2-multimer (v3) (Evans et al., 2021; Jumper et al., 2021). All structures generated using AlphaFold2-multimer (v3) are deposited to ZENODO (<u>10.5281/zenodo.10244735</u>). Four of the top-ranked structures resembled a V-shape dimer, with only the fourth-ranked structure displaying the experimentally known PI(4,5)P_2_ binding pocket facing towards a membrane-bound state. As the C95 residues were not in direct contact, we replaced the entire FGF2 dimer with the one obtained from atomistic MD simulations upon binding to the membrane. However, AlphaFold2-multimer (v3) predicted that the two Halo domains were too tightly bound together, which did not align well with the V-shape density resolved with cryo-ET. To address this, we used the “Fit in Map” command provided by the ChimeraX software to locally optimize the fit of one of the two Halo domains’ atomic coordinates into the density map (Goddard et al., 2018; Pettersen et al., 2021). As a result, we ensured no clashes with the FGF2 dimer and then applied the exact translation and rotation to the other Halo domain to create a symmetric structure. Finally, we positioned the modeled V-shape C95-C95 disulfide-bridged FGF2 dimer on the surface of a model membrane [POPC: CHOL (70:30) with 2 PI(4,5)P_2_ molecules] and subjected it to a 500 nanosecond simulation. We conducted three independent repeats and generated diversity by assigning random initial velocities. The topology of the V-shape C95 disulfide-bridged dimer was generated using the CHARMM-GUI website.

### Cryo-electron tomography and subtomogram averaging

Proteoliposomes containing FGF2-Y81pCMF-Halo (20 µl) were supplemented with 2.7 µl of Protein A-colloidal gold (10 nm) and applied onto glow discharged Quantifoil grids (R3.5/1). Glow discharging was done using Gatan Solarus 950. Proteoliposomes were vitrified by plunge freezing into liquid ethane using Vitrobot Mark IV (Thermo Fischer Scientific) with the following experimental settings: volume 3.5 µl, temperature 6 °C, humidity 100%, blotting force 0, 3-second blotting time). Cryo-ET was performed using the Titan Krios (ThermoFisher Scientific) equipped with an energy filter and K2 direct electron detector (Gatan). Tilt series were acquired at a magnification of 64,000x either in-focus using Volta phase plate (VPP) at EMBL, Heidelberg (pixel size 0.213 nm) or at nominal defocus −4 µm without the VPP at Heidelberg University (pixel size 0.229 nm) using an energy filter zero loss peak window set to 20eV. Mapping and tilt series acquisition was done using SerialEM (Mastronarde, 2005) and dose-symmetric tilt series schema with an increment angle 3° and a tilt range of 60° with cumulative electron dose 140 e^-^/Å^2^ (Hagen et al., 2017). Projection records were collected in counting mode as dose-fractionated movies and aligned on-fly using SerialEM 4K plug-in. Tilt series alignment with gold fiducial markers, CTF correction (except for VPP data), dose-filtration and tomogram reconstruction were done in IMOD (etomo GUI) using weighted back projection with SIRT-like filter with 7 iterations (Mastronarde and Held, 2017). Subtomogram extraction and averaging were performed in Dynamo (Castano-Diez et al., 2012; Castano-Diez, 2017; Castano-Diez et al., 2017; Navarro et al., 2018). At first, V-shaped particles were selected (N=186) using a dipole model with a north direction normal to the centre of the liposome at different positions with respect to the tilt axis of the tomogram acquired with VPP. Subvolumes with a cubic size of 160 voxels were extracted and averaged to create a template. Particles were iteratively aligned and averaged using a mask focusing on the protein density and a membrane and alternating c1 and c2 symmetry at each iteration. The final subtomogram average was visualized in ChimeraX (Goddard et al., 2018; Pettersen et al., 2021) where manual docking of Halo domain (PDB:4KAJ) and FGF2 monomer (PDB:1BFF) crystal structures was performed.

## Data availability

All structures generated using AlphaFold2-multimer (v3), as well as all initial and configuration molecular dynamics simulations files related to this study, have been deposited to ZENODO (<u>10.5281/zenodo.10244735</u>). The machine learning code to analyze MD simulations will be made publicly available upon separate publication. Detailed information regarding the repository location and licensing will be provided upon publication. Please contact Prof. Ilpo Vattulainen for further inquiries or access requests before public release. The source data underpinning the plots presented in this study are available for download as supplementary Excel files:

## Source files

**Figure 1—source data 1**

Real-time single-molecule TIRF recruitment assay.

**Figure 1—source data 2**

Cell surface biotinylation assay.

**Figure 2—source data 1**

Oligomeric size distribution of FGF2-GFP variants.

**Figure 3—source data 1**

Cross-linking quantification of FGF2 dimer to FGF2 monomer ratios.

**Figure 4—source data 1**

Membrane pore formation triggered by FGF2 oligomers.

**Figure 5—source data 1**

FGF2 membrane translocation and membrane pore formation assay (PI(4,5)P2 containing liposomes).

**Figure 6—source data 1**

FGF2 membrane translocation and membrane pore formation assay (Ni-NTA containing liposomes).

**Figure 7—source data 1**

Kinetic analysis of FGF2 / α1-subCD3 interaction.

**Figure 8—source data 1**

Populations of the individual clusters in the Bayesian Gaussian Mixture Model.

**Figure 9—source data 1**

Umbrella sampling free energy profile.

**Figure 10—source data 1**

Cross-Linking Mass Spectrometry fragments.

**Figure 11–video supplement 1.**

Simulation of the modeled V-Shape dimer interacting with the membrane surface over 500 nanoseconds.

## Acknowledgements

This work was supported by the Deutsche Forschungsgemeinschaft (SFB/TRR 186, project A1 and A5; WN and CF, DFG Ni 423/10-1, DFG Ni 423/12-1 and DFG Ni 423/13-1; WN and DFG LO 2821/1-1; FL). RŠ and MH acknowledge GAČR grant 20-01401J. We thank the cryo-EM network at Heidelberg University (HD-cryoNET) for support and assistance and the Electron Microscopy Core Facility at EMBL and Wim Hagen for data acquisition funded by iNext. The authors gratefully acknowledge the data storage service SDS@hd supported by the Ministry of Science, Research, and the Arts Baden-Württemberg (MWK), the German Research Foundation (DFG) through grant INST 35/1314-1 FUGG and INST 35/1503-1 FUGG. We also would like to acknowledge support from Holger Lorenz (ZMBH Imaging facility) and Monika Langlotz (ZMBH FACS Facility). The Vattulainen group has been supported by the Academy of Finland (projects 331349, 336234, 346135), the Sigrid Juselius Foundation, Helsinki Institute of Life Science (HiLIFE) Fellow Program, the Human Frontier Science Program (RGP0059/2019), the Lundbeck Foundation and DFG (SFB/TRR 83) (IV). This project has received funding from the European Union’s Horizon 2020 research and innovation programme under the Marie Skłodowska-Curie grant agreement No 101033606 (S.K.). We acknowledge the computing resources provided by the CSC – IT Center for Science Ltd. (Espoo, Finland). We further would like to thank Tobias P. Dick (DKFZ Heidelberg) who provided valuable insights into oxidative processes producing disulfide-bridged protein complexes at the inner plasma membrane leaflet.

## References

Abraham, M., T. Murtola, R. Schulz, S. Páll, J. Smith, B. Hess, and E. Lindahl. 2015. GROMACS: High performance molecular simulations through multi-level parallelism from laptops to supercomputers. SoftwareX. 1.

Akl, M.R., P. Nagpal, N.M. Ayoub, B. Tai, S.A. Prabhu, C.M. Capac, M. Gliksman, A. Goy, and K.S. Suh. 2016. Molecular and clinical significance of fibroblast growth factor 2 (FGF2 /bFGF) in malignancies of solid and hematological cancers for personalized therapies. Oncotarget. 7:44735–44762.

Albritton, L.M., L. Tseng, D. Scadden, and J.M. Cunningham. 1989. A putative murine ecotropic retrovirus receptor gene encodes a multiple membrane-spanning protein and confers susceptibility to virus infection. Cell. 57:659–666.

Amblard, I., E. Dupont, I. Alves, J. Miralves, I. Queguiner, and A. Joliot. 2020. Bidirectional transfer of homeoprotein EN2 across the plasma membrane requires PIP2. J Cell Sci. 133.

Backhaus, R., C. Zehe, S. Wegehingel, A. Kehlenbach, B. Schwappach, and W. Nickel. 2004. Unconventional protein secretion: membrane translocation of FGF-2 does not require protein unfolding. J Cell Sci. 117:1727–1736.

Benda, A., M. Benes, V. Marecek, A. Lhotsky, W. Hermens, and M. Hof. 2003. How To Determine Diffusion Coefficients in Planar Phospholipid Systems by Confocal Fluorescence Correlation Spectroscopy. Langmuir. 19:4120–4126.

Brown, D.I., and K.K. Griendling. 2009. Nox proteins in signal transduction. Free Radic Biol Med. 47:1239–1253.

Castano-Diez, D. 2017. The Dynamo package for tomography and subtomogram averaging: components for MATLAB, GPU computing and EC2 Amazon Web Services. Acta Crystallogr D Struct Biol. 73:478–487

Castano-Diez, D., M. Kudryashev, M. Arheit, and H. Stahlberg. 2012. Dynamo: a flexible, user-friendly development tool for subtomogram averaging of cryo-EM data in high-performance computing environments. J Struct Biol. 178:139–151.

Castano-Diez, D., M. Kudryashev, and H. Stahlberg. 2017. Dynamo Catalogue: Geometrical tools and data management for particle picking in subtomogram averaging of cryo-electron tomograms. J Struct Biol. 197:135–144.

Chaudhury, S., and J.J. Gray. 2008. Conformer selection and induced fit in flexible backbone protein-protein docking using computational and NMR ensembles. J Mol Biol. 381:1068–1087.

Chiritoiu, M., N. Brouwers, G. Turacchio, M. Pirozzi, and V. Malhotra. 2019. GRASP55 and UPR Control Interleukin-1beta Aggregation and Secretion. Developmental cell. 49:145–155 e144.

Dahl, J.P., A. Binda, V.A. Canfield, and R. Levenson. 2000. Participation of Na,K-ATPase in FGF-2 Secretion: Rescue of Ouabain-Inhibitable FGF-2 Secretion by Ouabain-Resistant Na,K-ATPase alpha Subunits. Biochemistry. 39:14877–14883.

Darden, T., D. York, and L. Pedersen. 1993. Particle mesh Ewald: An N-log(N) method for Ewald sums in large systems. J. Chem. Phys. 98:10089–10092.

Daura, X., K. Gademann, B. Jaun, D. Seebach, W. van Gunsteren, and A. Mark. 1999. Peptide folding: Ehen simulation meets experiment. Angew Chem Int Ed Engl. 38:236–240.

Debaisieux, S., F. Rayne, H. Yezid, and B. Beaumelle. 2012. The ins and outs of HIV-1 Tat. Traffic. 13:355–363.

Decker, C.G., Y. Wang, S.J. Paluck, L. Shen, J.A. Loo, A.J. Levine, L.S. Miller, and H.D. Maynard. 2016. Fibroblast growth factor 2 dimer with superagonist in vitro activity improves granulation tissue formation during wound healing. Biomaterials. 81:157–168.

Dimou, E., K. Cosentino, E. Platonova, U. Ros, M. Sadeghi, P. Kashyap, T. Katsinelos, S. Wegehingel, F. Noe, A.J. Garcia-Saez, H. Ewers, and W. Nickel. 2019. Single event visualization of unconventional secretion of FGF2. J Cell Biol. 218:683–699.

Dimou, E., and W. Nickel. 2018. Unconventional mechanisms of eukaryotic protein secretion. Curr Biol. 28:R406–R410.

Dupont, N., S. Jiang, M. Pilli, W. Ornatowski, D. Bhattacharya, and V. Deretic. 2011. Autophagy-based unconventional secretory pathway for extracellular delivery of IL-1beta. Embo J. 30:4701–4711.

Ebert, A.D., M. Laussmann, S. Wegehingel, L. Kaderali, H. Erfle, J. Reichert, J. Lechner, H.D. Beer, R. Pepperkok, and W. Nickel. 2010. Tec-kinase-mediated phosphorylation of fibroblast growth factor 2 is essential for unconventional secretion. Traffic. 11:813–826.

Engling, A., R. Backhaus, C. Stegmayer, C. Zehe, C. Seelenmeyer, A. Kehlenbach, B. Schwappach, S. Wegehingel, and W. Nickel. 2002. Biosynthetic FGF-2 is targeted to non-lipid raft microdomains following translocation to the extracellular surface of CHO cells. J. Cell Sci. 115:3619–3631.

Evans, D.J., and B.L. Holian. 1985. The Nose–Hoover thermostat. J. Chem. Phys. 83:4096.

Evans, R., M. O’Neill, A. Pritzel, N. Antropova, A. Senior, T. Green, A. Žídek, R. Bates, S. Blackwell, J. Yim, O. Ronneberger, S. Bodenstein, M. Zielinski, A. Bridgland, A. Potapenko, A. Cowie, K. Tunyasuvunakool, R. Jain, E. Clancy, P. Kohli, J. Jumper, and D. Hassabis. 2021. Protein complex prediction with AlphaFold-Multimer. Bioinformatics.

Evavold, C.L., J. Ruan, Y. Tan, S. Xia, H. Wu, and J.C. Kagan. 2017. The Pore-Forming Protein Gasdermin D Regulates Interleukin-1 Secretion from Living Macrophages. Immunity.

Florkiewicz, R.Z., J. Anchin, and A. Baird. 1998. The inhibition of fibroblast growth factor-2 export by cardenolides implies a novel function for the catalytic subunit of Na+,K+-ATPase. J Biol Chem. 273:544–551.

Goddard, T.D., C.C. Huang, E.C. Meng, E.F. Pettersen, G.S. Couch, J.H. Morris, and T.E. Ferrin. 2018. UCSF ChimeraX: Meeting modern challenges in visualization and analysis. Protein Sci. 27:14–25.

Gray, J.J., S. Moughon, C. Wang, O. Schueler-Furman, B. Kuhlman, C.A. Rohl, and D. Baker. 2003. Protein-protein docking with simultaneous optimization of rigid-body displacement and side-chain conformations. J Mol Biol. 331:281–299.

Hagen, W.J.H., W. Wan, and J.A.G. Briggs. 2017. Implementation of a cryo-electron tomography tilt-scheme optimized for high resolution subtomogram averaging. J Struct Biol. 197:191–198.

Hakim, M., and D. Fass. 2010. Cytosolic disulfide bond formation in cells infected with large nucleocytoplasmic DNA viruses. Antioxid Redox Signal. 13:1261–1271.

Heilig, R., M.S. Dick, L. Sborgi, E. Meunier, S. Hiller, and P. Broz. 2017. The Gasdermin-D pore acts as a conduit for IL-1beta secretion in mice. Eur J Immunol.

Huang, J., S. Rauscher, G. Nawrocki, T. Ran, M. Feig, B.L. de Groot, H. Grubmuller, and A.D. MacKerell, Jr. 2017. CHARMM36m: an improved force field for folded and intrinsically disordered proteins. Nat Methods. 14:71–73.

Iacobucci, C., C. Piotrowski, R. Aebersold, B.C. Amaral, P. Andrews, K. Bernfur, C. Borchers, N.I. Brodie, J.E. Bruce, Y. Cao, S. Chaignepain, J.D. Chavez, S. Claverol, J. Cox, T. Davis, G. Degliesposti, M.Q. Dong, N. Edinger, C. Emanuelsson, M. Gay, M. Gotze, F. Gomes-Neto, F.C. Gozzo, C. Gutierrez, C. Haupt, A.J.R. Heck, F. Herzog, L. Huang, M.R. Hoopmann, N. Kalisman, O. Klykov, Z. Kukacka, F. Liu, M.J. MacCoss, K. Mechtler, R. Mesika, R.L. Moritz, N. Nagaraj, V. Nesati, A.G.C. Neves-Ferreira, R. Ninnis, P. Novak, F.J. O’Reilly, M. Pelzing, E. Petrotchenko, L. Piersimoni, M. Plasencia, T. Pukala, K.D. Rand, J. Rappsilber, D. Reichmann, C. Sailer, C.P. Sarnowski, R.A. Scheltema, C. Schmidt, D.C. Schriemer, Y. Shi, J.M. Skehel, M. Slavin, F. Sobott, V. Solis-Mezarino, H. Stephanowitz, F. Stengel, C.E. Stieger, E. Trabjerg, M. Trnka, M. Vilaseca, R. Viner, Y. Xiang, S. Yilmaz, A. Zelter, D. Ziemianowicz, A. Leitner, and A. Sinz. 2019. First Community-Wide, Comparative Cross-Linking Mass Spectrometry Study. Anal Chem. 91:6953–6961.

Joliot, A., and A. Prochiantz. 2022. Unconventional Secretion, Gate to Homeoprotein Intercellular Transfer. Front Cell Dev Biol. 10:926421.

Jumper, J., R. Evans, A. Pritzel, T. Green, M. Figurnov, O. Ronneberger, K. Tunyasuvunakool, R. Bates, A. Zidek, A. Potapenko, A. Bridgland, C. Meyer, S.A.A. Kohl, A.J. Ballard, A. Cowie, B. Romera-Paredes, S. Nikolov, R. Jain, J. Adler, T. Back, S. Petersen, D. Reiman, E. Clancy, M. Zielinski, M. Steinegger, M. Pacholska, T. Berghammer, S. Bodenstein, D. Silver, O. Vinyals, A.W. Senior, K. Kavukcuoglu, P. Kohli, and D. Hassabis. 2021. Highly accurate protein structure prediction with AlphaFold. Nature. 596:583–589.

Kastrup, J.S., E.S. Eriksson, H. Dalboge, and H. Flodgaard. 1997. X-ray structure of the 154-amino-acid form of recombinant human basic fibroblast growth factor. comparison with the truncated 146-amino-acid form. Acta crystallographica. 53:160–168.

Katsinelos, T., W.A. McEwan, T.R. Jahn, and W. Nickel. 2021. Identification of cis-acting determinants mediating the unconventional secretion of tau. Scientific reports. 11:12946.

Katsinelos, T., M. Zeitler, E. Dimou, A. Karakatsani, H.M. Muller, E. Nachman, J.P. Steringer, C. Ruiz de Almodovar, W. Nickel, and T.R. Jahn. 2018. Unconventional Secretion Mediates the Trans-cellular Spreading of Tau. Cell Rep. 23:2039–2055.

La Venuta, G., S. Wegehingel, P. Sehr, H.M. Muller, E. Dimou, J.P. Steringer, M. Grotwinkel, N. Hentze, M.P. Mayer, D.W. Will, U. Uhrig, J.D. Lewis, and W. Nickel. 2016. Small Molecule Inhibitors Targeting Tec Kinase Block Unconventional Secretion of Fibroblast Growth Factor 2. J Biol Chem. 291:17787–17803.

La Venuta, G., M. Zeitler, J.P. Steringer, H.M. Müller, and W. Nickel. 2015. The Startling Properties of Fibroblast Growth Factor 2: How to Exit Mammalian Cells without a Signal Peptide at Hand. J Biol Chem. 290:27015–27020.

Lee, J., X. Cheng, J.M. Swails, M.S. Yeom, P.K. Eastman, J.A. Lemkul, S. Wei, J. Buckner, J.C. Jeong, Y. Qi, S. Jo, V.S. Pande, D.A. Case, C.L. Brooks, 3rd, A.D. MacKerell, Jr., J.B. Klauda, and W. Im. 2016. CHARMM-GUI Input Generator for NAMD, GROMACS, AMBER, OpenMM, and CHARMM/OpenMM Simulations Using the CHARMM36 Additive Force Field. J Chem Theory Comput. 12:405–413.

Legrand, C., R. Saleppico, J. Sticht, F. Lolicato, H.M. Muller, S. Wegehingel, E. Dimou, J.P. Steringer, H. Ewers, I. Vattulainen, C. Freund, and W. Nickel. 2020. The Na,K-ATPase acts upstream of phosphoinositide PI(4,5)P2 facilitating unconventional secretion of Fibroblast Growth Factor 2. Commun Biol. 3:141.

Lennicke, C., and H.M. Cocheme. 2021. Redox metabolism: ROS as specific molecular regulators of cell signaling and function. Mol Cell. 81:3691–3707.

Liu, F., P. Lossl, R. Scheltema, R. Viner, and A.J.R. Heck. 2017. Optimized fragmentation schemes and data analysis strategies for proteome-wide cross-link identification. Nat Commun. 8:15473.

Liu, L., M. Zhang, and L. Ge. 2020. Protein translocation into the ERGIC: an upstream event of secretory autophagy. Autophagy:1–3.

Liu, X., Z. Zhang, J. Ruan, Y. Pan, V.G. Magupalli, H. Wu, and J. Lieberman. 2016. Inflammasome-activated gasdermin D causes pyroptosis by forming membrane pores. Nature. 535:153–158.

Locker, J.K., and G. Griffiths. 1999. An unconventional role for cytoplasmic disulfide bonds in vaccinia virus proteins. J Cell Biol. 144:267–279.

Lolicato, F., and W. Nickel. 2022. A Role for Liquid-Ordered Plasma Membrane Nanodomains Coordinating the Unconventional Secretory Pathway of Fibroblast Growth Factor 2? Front Cell Dev Biol. 10:864257.

Lolicato, F., R. Saleppico, A. Griffo, A. Meyer, F. Scollo, B. Pokrandt, H.M. Muller, H. Ewers, H. Hahl, J.B. Fleury, R. Seemann, M. Hof, B. Brugger, K. Jacobs, I. Vattulainen, and W. Nickel. 2022. Cholesterol promotes clustering of PI(4,5)P2 driving unconventional secretion of FGF2. J Cell Biol. 221.

Malhotra, V. 2013. Unconventional protein secretion: an evolving mechanism. EMBO J. 32:1660–1664.

Mastronarde, D.N. 2005. Automated electron microscope tomography using robust prediction of specimen movements. J Struct Biol. 152:36–51.

Mastronarde, D.N., and S.R. Held. 2017. Automated tilt series alignment and tomographic reconstruction in IMOD. J Struct Biol. 197:102–113.

Meng, Y., and B. Roux. 2015. Efficient Determination of Free Energy Landscapes in Multiple Dimensions from Biased Umbrella Sampling Simulations Using Linear Regression. J Chem Theory Comput. 11:3523–3529.

Merezhko, M., C.A. Brunello, X. Yan, H. Vihinen, E. Jokitalo, R.L. Uronen, and H.J. Huttunen. 2018. Secretion of Tau via an Unconventional Non-vesicular Mechanism. Cell Rep. 25:2027–2035 e2024.

Merezhko, M., R.L. Uronen, and H.J. Huttunen. 2020. The Cell Biology of Tau Secretion. Front Mol Neurosci. 13:569818.

Mirdita, M., K. Schutze, Y. Moriwaki, L. Heo, S. Ovchinnikov, and M. Steinegger. 2022. ColabFold: making protein folding accessible to all. Nat Methods. 19:679–682.

Monteleone, M., A.C. Stanley, K.W. Chen, D.L. Brown, J.S. Bezbradica, J.B. von Pein, C.L. Holley, D. Boucher, M.R. Shakespear, R. Kapetanovic, V. Rolfes, M.J. Sweet, J.L. Stow, and K. Schroder. 2018. Interleukin-1beta Maturation Triggers Its Relocation to the Plasma Membrane for Gasdermin-D-Dependent and - Independent Secretion. Cell Rep. 24:1425–1433.

Müller, H.M., J.P. Steringer, S. Wegehingel, S. Bleicken, M. Munster, E. Dimou, S. Unger, G. Weidmann, H. Andreas, A.J. Garcia-Saez, K. Wild, I. Sinning, and W. Nickel. 2015. Formation of Disulfide Bridges Drives Oligomerization, Membrane Pore Formation and Translocation of Fibroblast Growth Factor 2 to Cell Surfaces. J Biol Chem. 290:8925–8937.

Navarro, P.P., H. Stahlberg, and D. Castano-Diez. 2018. Protocols for Subtomogram Averaging of Membrane Proteins in the Dynamo Software Package. Front Mol Biosci. 5:82.

Nawrocka, D., M.A. Krzyscik, L. Opalinski, M. Zakrzewska, and J. Otlewski. 2020. Stable Fibroblast Growth Factor 2 Dimers with High Pro-Survival and Mitogenic Potential. Int J Mol Sci. 21.

Netto, L.E.S., and L. Machado. 2022. Preferential redox regulation of cysteine-based protein tyrosine phosphatases: structural and biochemical diversity. FEBS J. 289:5480–5504.

Nickel, W. 2007. Unconventional secretion: an extracellular trap for export of fibroblast growth factor 2. J Cell Sci. 120:2295–2299.

Nickel, W. 2011. The unconventional secretory machinery of fibroblast growth factor 2. Traffic. 12:799–805.

Nickel, W., and C. Rabouille. 2009. Mechanisms of regulated unconventional protein secretion. Nat Rev Mol Cell Biol. 10:148–155.

Nickel, W., and M. Seedorf. 2008. Unconventional mechanisms of protein transport to the cell surface of eukaryotic cells. Annu Rev Cell Dev Biol. 24:287–308.

Nordzieke, D.E., and I. Medrano-Fernandez. 2018. The Plasma Membrane: A Platform for Intra- and Intercellular Redox Signaling. Antioxidants (Basel). 7.

Pallotta, M.T., and W. Nickel. 2020. FGF2 and IL-1beta - explorers of unconventional secretory pathways at a glance. J Cell Sci. 133.

Parrinello, M., and A. Rahman. 1981. Polymorphic transitions in single crystals: A new molecular dynamics method. J. Appl. Phys. 52:7182–7190.

Paszke, A., S. Gross, F. Massa, A. Lerer, J. Bradbury, G. Chanan, T. Killeen, Z. Lin, N. Gimelshein, L. Antiga, A. Desmaison, A. Kopf, E. Yang, Z. DeVito, M. Raison, A. Tejani, S. Chilamkurthy, B. Steiner, L. Fang, J. Bai, and S. Chintala. 2019. PyTorch: An Imperative Style, High-Performance Deep Learning Library. In Advances in Neural Information Processing Systems 32. Curran Associates, Inc. 8024–8035.

Pedregosa, F., G. Varoquaux, A. Gramfort, V. Michel, B. Thirion, O. Grisel, M. Blondel, P. Prettenhofer, R. Weiss, V. Dubourg, J. Vanderplas, A. Passos, D. Cournapeau, M. Brucher, M. Perrot, and E. Duchesnay. 2011. Scikit-learn: Machine Learning in Python. Journal of Machine Learning Research. 12:2825–2830.

Pettersen, E.F., T.D. Goddard, C.C. Huang, E.C. Meng, G.S. Couch, T.I. Croll, J.H. Morris, and T.E. Ferrin. 2021. UCSF ChimeraX: Structure visualization for researchers, educators, and developers. Protein Sci. 30:70–82.

Piersimoni, L., and A. Sinz. 2020. Cross-linking/mass spectrometry at the crossroads. Anal Bioanal Chem. 412:5981–5987.

Plotnikov, A.N., S.R. Hubbard, J. Schlessinger, and M. Mohammadi. 2000. Crystal structures of two FGF-FGFR complexes reveal the determinants of ligand-receptor specificity. Cell. 101:413–424.

Plotnikov, A.N., J. Schlessinger, S.R. Hubbard, and M. Mohammadi. 1999. Structural basis for FGF receptor dimerization and activation. Cell. 98:641–650.

Rabouille, C. 2017. Pathways of Unconventional Protein Secretion. Trends Cell Biol. 27:230–240.

Rappsilber, J., M. Mann, and Y. Ishihama. 2007. Protocol for micro-purification, enrichment, pre-fractionation and storage of peptides for proteomics using StageTips. Nat Protoc. 2:1896–1906.

Rayne, F., S. Debaisieux, H. Yezid, Y.L. Lin, C. Mettling, K. Konate, N. Chazal, S.T. Arold, M. Pugniere, F. Sanchez, A. Bonhoure, L. Briant, E. Loret, C. Roy, and B. Beaumelle. 2010. Phosphatidylinositol-(4,5)-bisphosphate enables efficient secretion of HIV-1 Tat by infected T-cells. EMBO J. 29:1348–1362.

Rubin, D. 1981. The Bayesian Bootstrap. Ann. Statist. 9:130–134.

Sachl, R., S. Cujova, V. Singh, P. Riegerova, P. Kapusta, H.M. Muller, J.P. Steringer, M. Hof, and W. Nickel. 2020. Functional Assay to Correlate Protein Oligomerization States with Membrane Pore Formation. Anal Chem. 92:14861–14866.

Schäfer, T., H. Zentgraf, C. Zehe, B. Brugger, J. Bernhagen, and W. Nickel. 2004. Unconventional secretion of fibroblast growth factor 2 is mediated by direct translocation across the plasma membrane of mammalian cells. J Biol Chem. 279:6244–6251.

Schatz, M., P.B.V. Tong, and B. Beaumelle. 2018. Unconventional secretion of viral proteins. Seminars in cell & developmental biology. 83:8–11.

Schindelin, J., I. Arganda-Carreras, E. Frise, V. Kaynig, M. Longair, T. Pietzsch, S. Preibisch, C. Rueden, S. Saalfeld, B. Schmid, J.Y. Tinevez, D.J. White, V. Hartenstein, K. Eliceiri, P. Tomancak, and A. Cardona. 2012. Fiji: an open-source platform for biological-image analysis. Nat Methods. 9:676–682.

Schlessinger, J., A.N. Plotnikov, O.A. Ibrahimi, A.V. Eliseenkova, B.K. Yeh, A. Yayon, R.J. Linhardt, and M. Mohammadi. 2000. Crystal structure of a ternary FGF-FGFR-heparin complex reveals a dual role for heparin in FGFR binding and dimerization. Mol Cell. 6:743–750.

Seelenmeyer, C., S. Wegehingel, I. Tews, M. Kunzler, M. Aebi, and W. Nickel. 2005. Cell surface counter receptors are essential components of the unconventional export machinery of galectin-1. J Cell Biol. 171:373–381.

Sitia, R., and A. Rubartelli. 2018. The unconventional secretion of IL-1beta: Handling a dangerous weapon to optimize inflammatory responses. Seminars in cell & developmental biology. 83:12–21.

Sparn, C., E. Dimou, A. Meyer, R. Saleppico, S. Wegehingel, M. Gerstner, S. Klaus, H. Ewers, and W. Nickel. 2022a. Glypican-1 drives unconventional secretion of fibroblast growth factor 2. Elife. 11.

Sparn, C., A. Meyer, R. Saleppico, and W. Nickel. 2022b. Unconventional secretion mediated by direct protein self-translocation across the plasma membranes of mammalian cells. Trends Biochem Sci.

Steringer, J.P., S. Bleicken, H. Andreas, S. Zacherl, M. Laussmann, K. Temmerman, F.X. Contreras, T.A. Bharat, J. Lechner, H.M. Müller, J.A. Briggs, A.J. Garcia-Saez, and W. Nickel. 2012. Phosphatidylinositol 4,5-Bisphosphate (PI(4,5)P2)-dependent Oligomerization of Fibroblast Growth Factor 2 (FGF2) Triggers the Formation of a Lipidic Membrane Pore Implicated in Unconventional Secretion. J Biol Chem. 287:27659–27669.

Steringer, J.P., S. Lange, S. Cujova, R. Sachl, C. Poojari, F. Lolicato, O. Beutel, H.M. Muller, S. Unger, U. Coskun, A. Honigmann, I. Vattulainen, M. Hof, C. Freund, and W. Nickel. 2017. Key steps in unconventional secretion of fibroblast growth factor 2 reconstituted with purified components. Elife. 6:e28985.

Steringer, J.P., and W. Nickel. 2018. A direct gateway into the extracellular space: Unconventional secretion of FGF2 through self-sustained plasma membrane pores. Seminars in cell & developmental biology.

Temmerman, K., A.D. Ebert, H.M. Müller, I. Sinning, I. Tews, and W. Nickel. 2008. A direct role for phosphatidylinositol-4,5-bisphosphate in unconventional secretion of fibroblast growth factor 2. Traffic. 9:1204–1217.

Temmerman, K., and W. Nickel. 2009. A novel flow cytometric assay to quantify interactions between proteins and membrane lipids. J Lipid Res. 50:1245–1254.

Tinevez, J.Y., N. Perry, J. Schindelin, G.M. Hoopes, G.D. Reynolds, E. Laplantine, S.Y. Bednarek, S.L. Shorte, and K.W. Eliceiri. 2017. TrackMate: An open and extensible platform for single-particle tracking. Methods. 115:80–90.

Torrie, G., and V. JP. 1974. Monte Carlo free energy estimates using non-Boltzmann sampling: Application to the sub-critical Lennard-Jones fluid. Chemical Physics Letters. 28:578–581.

Torrie, G., and J. Valleau. 1977. Nonphysical sampling distributions in Monte Carlo free-energy estimation: Umbrella sampling. Journal of Computational Physics. 23:187–199.

Urlinger, S., U. Baron, M. Thellmann, M.T. Hasan, H. Bujard, and W. Hillen. 2000. Exploring the sequence space for tetracycline-dependent transcriptional activators: novel mutations yield expanded range and sensitivity. Proc. Natl. Acad. Sci. U.S.A. 97:7963–7968.

Verlet, L. 1967. Computer “Experiments” on Classical Fluids. I. Thermodynamical Properties of Lennard-Jones Molecules. Physical Review Journals Archive. 159:98–103.

Wang, C., P. Bradley, and D. Baker. 2007. Protein-protein docking with backbone flexibility. J Mol Biol. 373:503–519.

Wang, C., O. Schueler-Furman, and D. Baker. 2005. Improved side-chain modeling for protein-protein docking. Protein Sci. 14:1328–1339.

Wang, W., D. Yang, F. Chen, Y. Pang, S. Huang, and Y. Ge. 2019. Clustering With Orthogonal AutoEncoder. IEEE Access. 7:62421–62432.

Widengren, J., U. Mets, and R. Rigler. 1995. Fluorescence correlation spectroscopy of triplet states in solution: a theoretical and experimental study. J. Phys. Chem. 99:13368–13379.

Ye, Y. 2018. Regulation of protein homeostasis by unconventional protein secretion in mammalian cells. Seminars in cell & developmental biology. 83:29–35.

Zacherl, S., G. La Venuta, H.M. Müller, S. Wegehingel, E. Dimou, P. Sehr, J.D. Lewis, H. Erfle, R. Pepperkok, and W. Nickel. 2015. A direct role for ATP1A1 in unconventional secretion of fibroblast growth factor 2. J Biol Chem. 290:3654–3665.

Zehe, C., A. Engling, S. Wegehingel, T. Schafer, and W. Nickel. 2006. Cell-surface heparan sulfate proteoglycans are essential components of the unconventional export machinery of FGF-2. Proc Natl Acad Sci U S A. 103:15479–15484.

Zhang, M., L. Liu, X. Lin, Y. Wang, Y. Li, Q. Guo, S. Li, Y. Sun, X. Tao, D. Zhang, X. Lv, L. Zheng, and L. Ge. 2020. A Translocation Pathway for Vesicle-Mediated Unconventional Protein Secretion. Cell. 181:637–652 e615.

Zhang, M., and R. Schekman. 2013. Cell biology. Unconventional secretion, unconventional solutions. Science. 340:559–561.

