## Supplementary figures and images for "Disulfide bridge-dependent dimerization triggers FGF2 membrane translocation into the extracellular space"

### Supplemental Data 2

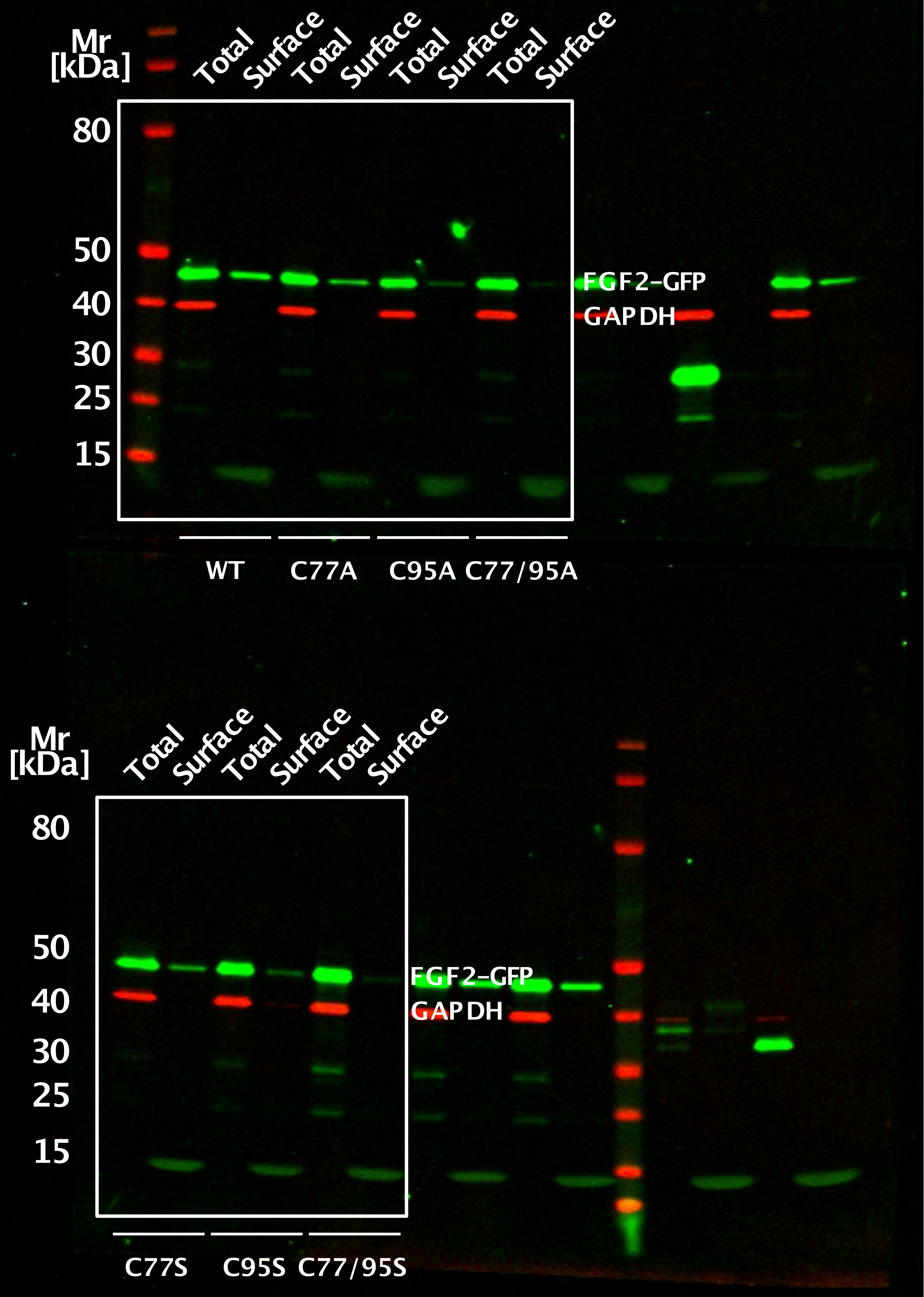
